# Complex spike firing adapts to saliency of inputs and engages readiness to act

**DOI:** 10.1101/2020.09.26.314534

**Authors:** Lorenzo Bina, Vincenzo Romano, Tycho M. Hoogland, Laurens W.J. Bosman, Chris I. De Zeeuw

**Author notes:** Correspondence: Laurens Bosman, or Chris De Zeeuw, Department of Neuroscience, PO Box 2040, 3000 CA Rotterdam, the Netherlands.

## Abstract

The cerebellum is involved in cognition next to motor coordination. During complex tasks, climbing fiber input to the cerebellum can deliver seemingly opposite signals, covering both motor and non-motor functions. To elucidate this ambiguity, we hypothesized that climbing fiber activity represents the saliency of inputs leading to action-readiness. We addressed this hypothesis by recording Purkinje cell activity in lateral cerebellum of awake mice learning go/no-go decisions based on entrained saliency of different sensory stimuli. As training progressed, the timing of climbing fiber signals switched in a coordinated fashion with that of Purkinje cell simple spikes towards the moment of occurrence of the salient stimulus that required action. Trial-by-trial analysis indicated that emerging climbing fiber activity is not linked to individual motor responses or rewards per se, but rather reflects the saliency of a particular sensory stimulus that engages a general readiness to act, bridging the non-motor with the motor functions.

**In brief:** 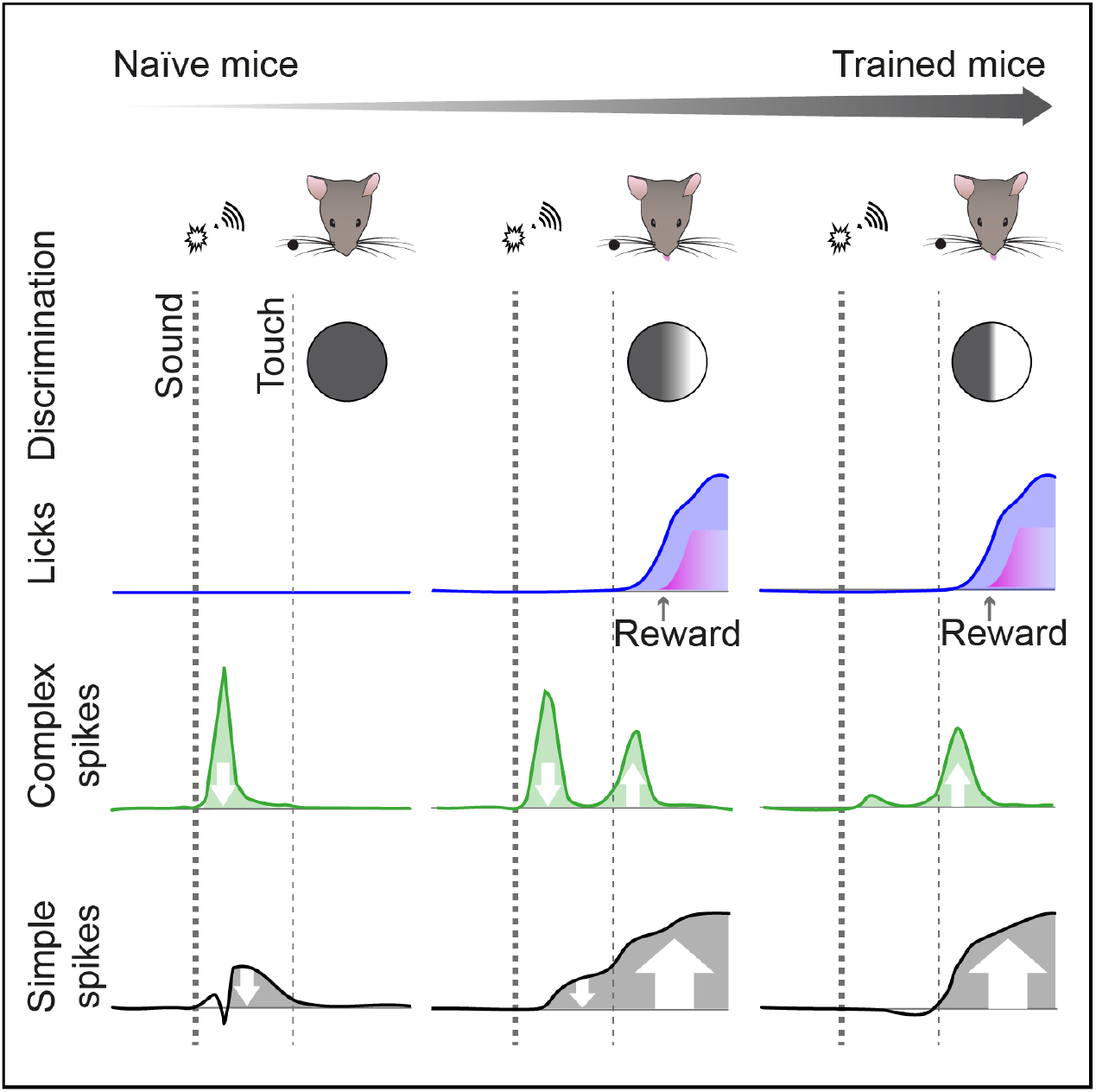

Mice were trained to identify the saliency of different sensory inputs in that they had to learn to ignore a prominent sound cue and respond to a light tactile cue in a Go/No-Go licking task. As the mice learned to discriminate the two inputs and respond to the proper signal, the Purkinje cells in the lateral cerebellum switched their climbing fiber activity (i.e., complex spike activity) towards the moment of occurrence of the salient stimulus that required a response, while concomitantly shifting the phase of their simple spike modulation. Trial-by-trial analysis indicates that the emerging climbing fiber activity is not linked to the occurrence of the motor response or reward per se, but rather reflects the saliency of a particular sensory stimulus engaging a general readiness to act.

## Introduction

The brain receives a continuous stream of sensory input, most of which can be ignored. The selection of salient information that require attention, however, can be a matter of life and death: ignoring the presence of a predator can easily be a fatal mistake. The saliency of sensory input depends on the behavioral and environmental context of an animal, and, as a consequence, the behavioral action taken in response to the same inputs can vary over time. This is true for relatively simple behaviors, such as the adaptation of the gill withdrawal reflex in sea snails that changes upon repeated touch (Castellucci et al., 1970; Frost et al., 1985), but also for more complex voluntary behaviors, such as ignoring a red sign when rushing to the hospital. Selective attention can even be a social phenomenon, as for instance sentinel behavior in meerkats involves the distribution of attention over group members (Clutton-Brock et al., 1999; Santema and Clutton-Brock, 2013).

Selective attention is closely related to the working memory and often considered to be organized by the forebrain in conjunction with the midbrain (Buschman and Kastner, 2015; Knudsen, 2018; Smith and Jonides, 1999). In contrast, the cerebellum is deemed crucial for the context-dependent adaptation of reflexes (Ito, 2000; Jirenhed et al., 2007; McCormick and Thompson, 1984; Romano et al., 2018; Ten Brinke et al., 2015). While the cerebellum is known to be involved in the execution of voluntary and autonomic behavior (Boyd, 2010; Romano et al., 2020; Sauerbrei et al., 2015; Vinueza Veloz et al., 2015), it is unclear to what extent the cerebellum is required for selective attention in relation to voluntary behavior. On the one hand, cerebellar patients do not necessarily have attentional deficits (Helmuth et al., 1997) - although the interpretation of this finding can be obfuscated by a residual function of the cerebellum and/or compensation by other brain regions (Abdelgabar et al., 2019). On the other hand, human brain imaging does show cerebellar activity during focused as well as shifting attention, even in the absence of movements (Allen et al., 1997; Brissenden et al., 2018; Le et al., 1998).

According to classical theories, cerebellar Purkinje cells, which form the sole output of the cerebellar cortex, can adjust the weights of their sensory inputs through mechanisms of supervised learning (Albus, 1971; Marr, 1969). Purkinje cells generate high-frequent simple spikes and low-frequent complex spikes (Thach, 1967; Zhou et al., 2014). Whereas simple spike modulation is mediated by changes in activity of the mossy fibers, which originate from various sources in the brainstem, complex spike modulation results from changes in activity of the climbing fibers, all of which are derived from the inferior olive (De Zeeuw et al., 2011). Simple spike plasticity, under control of climbing fiber activity, can change the motor response to sensory feedback (Herzfeld et al., 2018; Ohmae and Medina, 2015; Romano et al., 2018; Ten Brinke et al., 2015; Yang and Lisberger, 2014). However, the role of cerebellar plasticity during tasks that include not only motor but also non-motor functions is still enigmatic (Chabrol et al., 2019; Deverett et al., 2018; Gao et al., 2018; Heffley et al., 2018; Hull, 2020; Kostadinov et al., 2019; Larry et al., 2019; Sendhilnathan et al., 2020; Tsutsumi et al., 2019). In particular, it is unclear when climbing fibers are activated during the acquisition of such tasks, to what extent they are causally linked to the related entrained movements as well as the expectation, presentation and/or omission of rewards, and what their impact is on concomitant simple spike modulation.

We hypothesized that climbing fiber modulation during complex, i.e., combined cognitive - motor, tasks may represent the saliency of particular sensory inputs, setting the stage for a behavioral response without actually encoding the triggered motor activity, just like the starter signals the onset of a race without directly controlling the athletes’ movements. In other words, we presumed that climbing fiber activity can be tuned to engage a readiness to act, allowing the animal to make the appropriate response based on a selection of salient sensory and/or internal signals. We addressed this hypothesis by studying Purkinje cell activity in lateral cerebellum of awake mice while they learned to make (i.e., decision to go) or to avoid (i.e., no-go) a licking movement based on the saliency of different sensory stimuli, i.e., a clear sound followed by a rod moving within (go cue) or outside (no-go cue) of their whisker field at a fixed temporal interval (Fig. 1A). Mice were trained to delay their response until they could base their decision on the presence or absence of the rod in their whisker field. During go trials, mice were rewarded with water when they licked during the response interval. When they licked too early, however, they were punished with an air puff and an extra delay till the next trial. We followed Purkinje cell activity with electrophysiological recordings and calcium-imaging throughout the learning process and show that Purkinje cells change their activity pattern, altering complex spike and simple spike patterns bidirectionally and reciprocally. Indeed, the sequence of complex spike responses and subsequent simple spike increases following the application of a particular stimulus could be transferred in time when the subject learned to recognize the more prominent saliency of another stimulus presented at a later stage of the task. Using optogenetic stimulation and genetic interference, we show that the changes in Purkinje cell activity enable effective learning of the sensory selection task, revealing a role for the climbing fibers in acquiring and mediating the saliency of input signals, while enhancing context-dependent readiness to act at the optimal moment. These findings highlight how climbing fiber signaling can set the pace when coordinating non-motor functions with motor responses.

**Figure 1.**
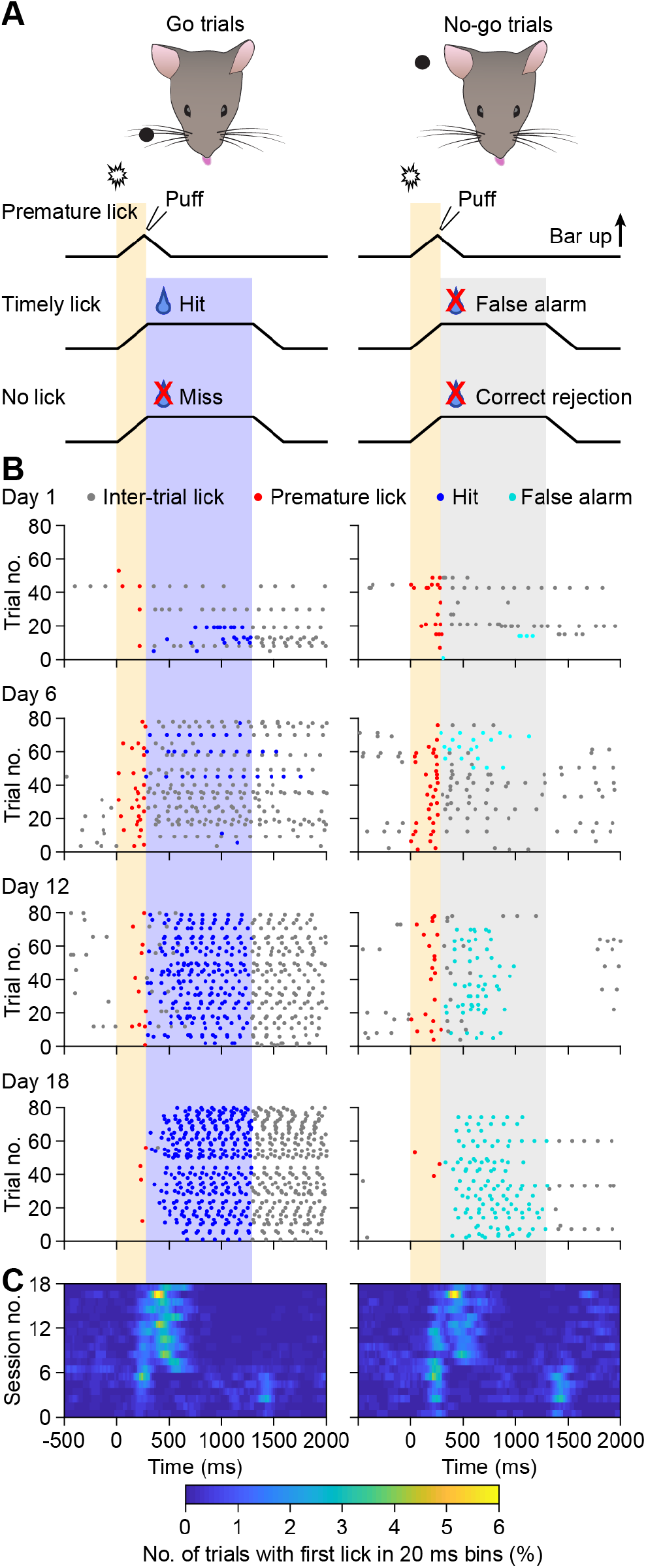
Training mice on a timed object detection task. **A**. Mice were trained on a timed object task in which they had to associate the position of a pole relative to their whisker field with the ability to obtain a water reward. In between trials, when the pole was well below the mice, the pole was rotated to be either below the whisker field or to a more posterior position: the former in preparation of go trials, the latter of no-go trials. At trial start, a clearly audible sound was made by the pneumatic valve controlling the rising of the pole. The onset of the sound was followed by a period of 300 ms during which the mice were not allowed to lick (yellow shade). Licking during this period triggered an aversive air puff to their nose and caused an immediate cessation of the trial. The mice then had to wait at least 7 s for the next trial to start. After the no-lick period, a response interval of 1000 ms followed (red / grey shades). During the response intervals of go trials, mice could activate the water valve to obtain a water reward, implying that water was only available after the mouse licked first. **B**. At training onset, mice typically started to engage in the task by licking. Initially they did not adhere to the different trial phases. During training, lick timing markedly improved. All raster plots come from the same mouse. **C**. Heat maps representing the averaged occurrences of first licks of bouts, showing first an increase in licking during the no-lick period, and afterwards a delay in the onset of lick bouts. The heat maps come from the same mouse as the data plotted in **B**.

## Results

### Training mice on a go/no-go paradigm with a no-response period

To what extent does plasticity of cerebellar Purkinje cell activity in the lateral cerebellum contribute to learning and well-timed execution of a sensory selection task? To answer this question, we trained head-fixed mice on a go/no-go task for 18 daily sessions. Every trial began with an auditory cue that was identical for go and no-go trials. The sound was created by a pneumatic valve that caused the rise of a metal pole into the whisker field. Around 300 ms later, the pole reached its maximal position, either within (go trials) or out of (no-go trials) reach of the facial whiskers. The mice were trained to suspend action for the 300 ms interval during which the pole rose into the whisker field, after which a response interval ensued of 1 s. During the response interval of go trials, licking triggered a water reward – the reward was not presented without prior action of the mouse. Licking during the 300 ms no-lick period induced an aversive air puff to the nose and caused an immediate cessation of the ongoing trial (Fig. 1A). Typically, mice first learned to engage in licking prior to withholding licking at undesired moments (Fig. 1B-C).

### Learning to suspend action takes longer than to act

Before the training started, mice were habituated to the setup and familiarized with the presence of the lick port. On the first day of training, mice did not yet fully engage with the task, showing only few trials with licks, but they became more engaged over the course of the first 8 days. Starting at on average day 5 of the training, mice began to differentiate between go and no-go trials, and showed a bias towards responding during go trials. During this same period, mice also licked more frequently during the no-response window, triggering early termination of trials. Typically around 8 days of training, mice began to withhold licking until the start of the response window (Fig. 2A-C). Mice often reached a plateau level of performance after two weeks of training without further improvements. Training was therefore stopped after 18 days, when the mice showed early licks during 26 ± 12% of the trials and correct behavior during 48 ± 11% of all trials (averages ± sd). As explained below, the relatively large fraction of error trials helped us to discriminate between neuronal correlates of decision making versus those of motor control.

**Figure 2.**
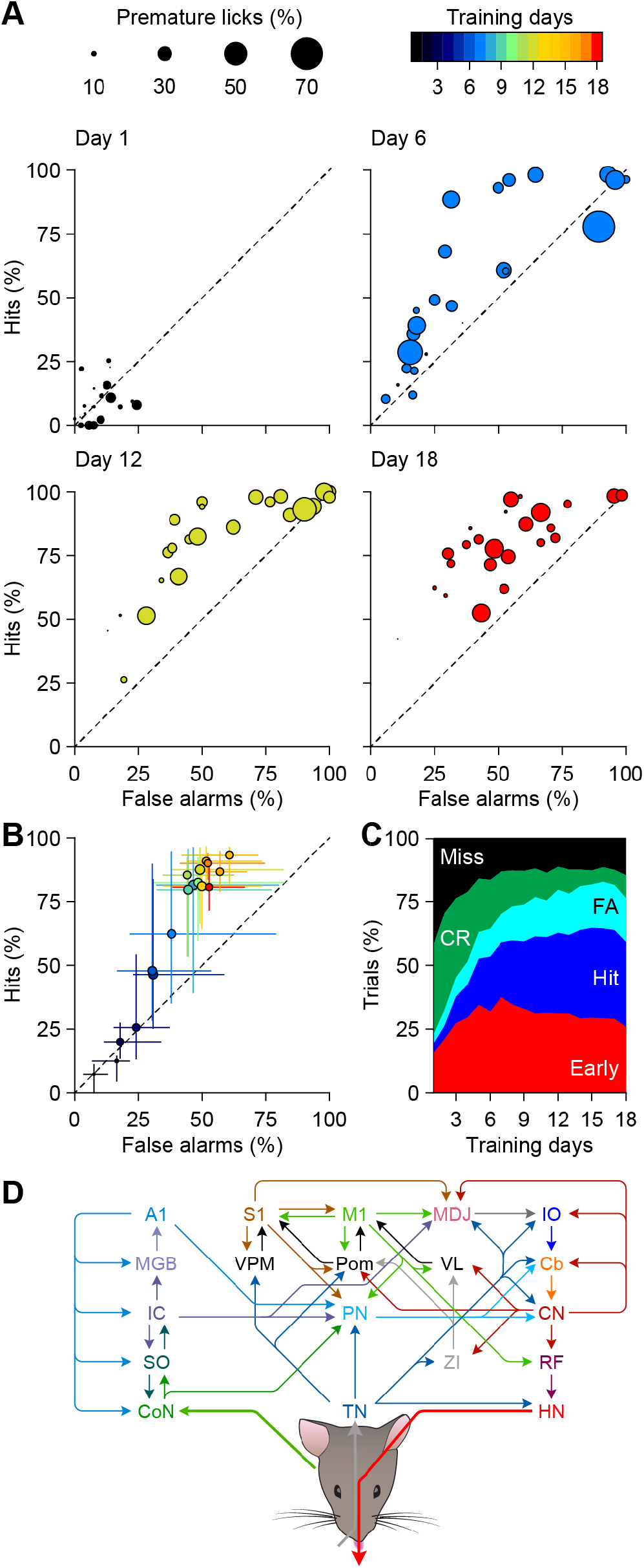
Learning to suspend licking takes more time than to learn to lick. **A**. Scatter plots of all 24 mice at four stages during training, comparing the fraction of trials with licks during go trials (y-axis) and no-go trials (x-axis). The diameter of the circles indicates the fraction of trials that were aborted due to premature licking. **B**. Median training performance of the 24 mice. Bars indicate inter-quartile range. Mice started in the lower left corner (no licking) and gradually moved first along the 45° line (more licking, but not discriminating between go and no-go trials), and later above the 45° line (more licking during go than during no-go trials). Days are indicated by color code (see top of **C**). **C**. Learning performance per trial type averaged over 24 mice. **D**. Simplified scheme of main anatomical pathways transporting auditory and whisker input and orchestrating tongue movements. A1 = primary auditory cortex; Cb = cerebellar cortex; CN = cerebellar nuclei; CoN = cochlear nucleus; HN = hypoglossal nucleus; IC = inferior colliculus; IO = inferior olive; M1 = primary motor cortex; MDJ = nuclei of the mesodiencephalic junction; PN = pontine nuclei; Pom = thalamic posteriomedial nucleus; RF = reticular formation; S1 = primary somatosensory cortex; SO = superior olive; TN = sensory trigeminal nuclei; VL = thalamic ventrolateral nucleus; VPM = thalamic ventral posteriomedial nucleus; ZI = zona incerta.

As our task required mice to make a well-timed response based on a combination of auditory and tactile inputs, a large number of brain regions is expected to be involved. On the sensory side, these include the auditory pathway, from the cochlear nucleus via the superior olive, the inferior colliculus and the medial geniculate body to the primary auditory cortex (A1), and the tactile pathway from the sensory trigeminal nuclei via the ventral posteromedial (VPM) and posteromedial (Pom) nuclei of the thalamus to the primary somatosensory cortex (S1) and the primary motor cortex (M1) (Bosman et al., 2011; Cant and Oliver, 2018; Moore, 1991; Yu et al., 2006). The motor output of the tongue is ultimately generated in the hypoglossal nucleus (Lowe, 1980; McElvain et al., 2018). In between are several cortical and subcortical structures, including the cerebellum with its central role in sensorimotor integration, timing and expectation (De Zeeuw et al., 2011; Ivry and Keele, 1989; Kostadinov et al., 2019; Moberget and Ivry, 2019; Rahmati et al., 2014). Purkinje cells of the lateral cerebellum receive auditory and whisker input via multiple pathways and can affect licking behavior indirectly via the reticular formation and hypoglossal nucleus (Borke et al., 1983; Bosman et al., 2011; Bosman et al., 2010; Ju et al., 2019; Steinmetz et al., 1987; Teune et al., 2000) (Fig. 2D).

### Distinct Purkinje cell responses of naïve and trained mice

As a first step towards understanding the interactions between Purkinje cell activity and task performance, we isolated sensory responses by recording from Purkinje cells in naïve mice. The naïve mice had been habituated to the recording setup, but were not accustomed to the lick port. As a consequence, none of them licked during the recording session. In the naïve mice, the sound cue that announced the start of the trials evoked a statistically significant complex spike response in 16 out of 24 (67%) recorded Purkinje cells. The complex spike response to the sound cue was stronger than that to the tactile stimulus, but did not differ significantly between go and no-go trials (*p* = 0.002 and *p* = 0.494, respectively, Dunn’s post-hoc tests after Friedman’s ANOVA, see Table S1 for more details on statistical analysis; Fig. 3A-C).

**Figure 3.**
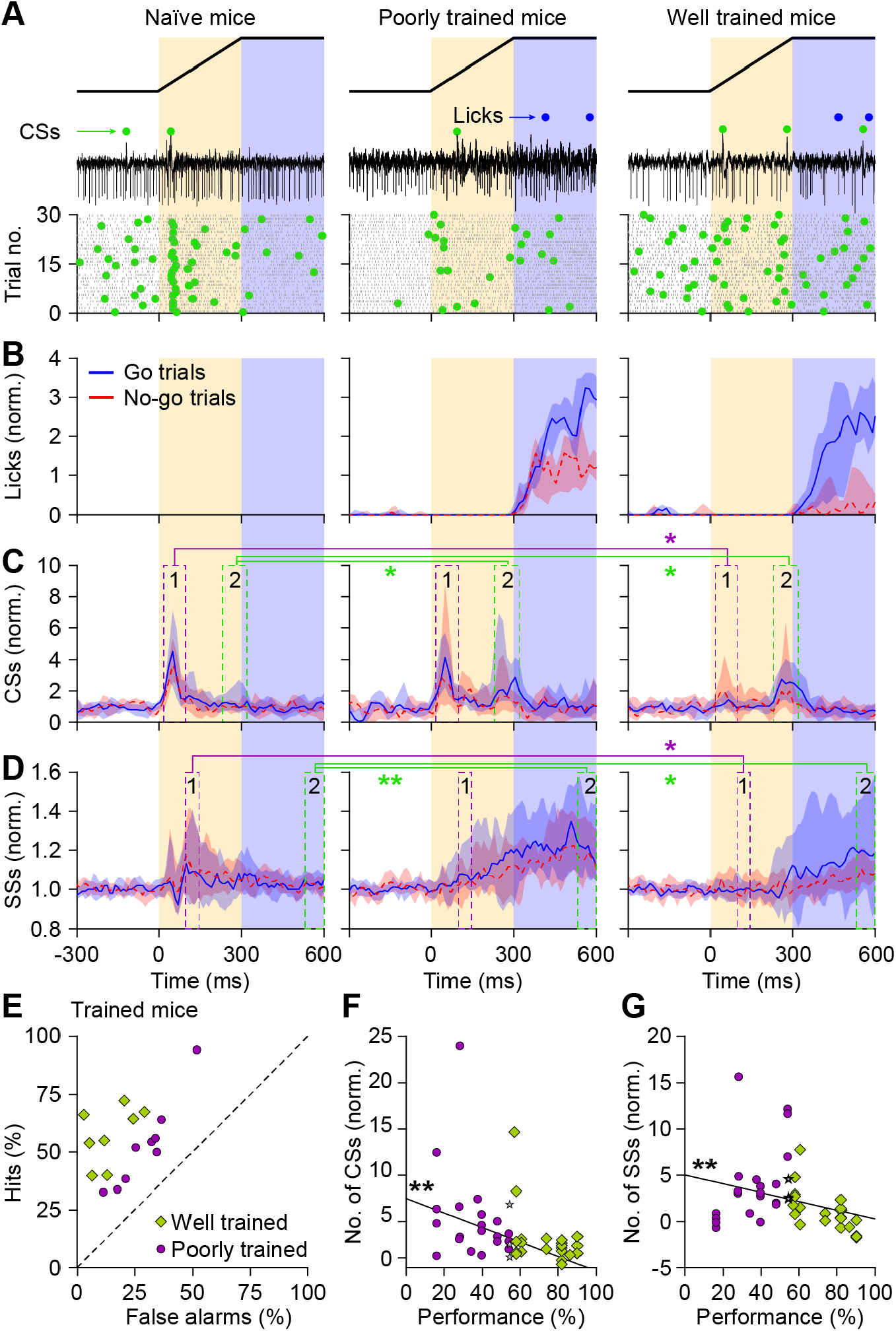
Distinct Purkinje cell responses in naïve and trained mice. **A**. Representative recordings of Purkinje cells in the crus 1 / crus 2 area of a naïve mouse (left), a trained mouse with a relatively poor performance (middle), and a trained mouse with a good performance (right). Purkinje cell recordings showed complex spikes (green dots) and simple spikes (black dots), with licks indicate above the trials with blue dots. **B**. Peri-stimulus time histograms of licks during go (blue) and no-go (red) trials in naïve, poorly trained and well-trained mice. Naïve mice were not aware of the ability to obtain water and consequently did not lick. Note that the poorly trained mice licked relatively often during no-go trials. **C**. Peri-stimulus time histograms of all Purkinje cells that showed a statistically significant modulation of their Purkinje cells (16, 15 and 12 cells per category, respectively). Relative to the naïve mice, the well-trained mice showed a decrease in the first (sound-evoked) and an increase in the second (touch-induced) complex spike peak, with the poorly trained mice showing more resemblance with naïve mice. **D**. Also simple spike modulation changed during training, reducing the early (sound-evoked) responses and increasing modulation during the response window. In **B, C** and **D**, the lines indicate the medians and the shaded area the inter-quartile range. In order to obtain a clearer comparison, trials terminated because of licking during the no-lick period have been excluded from this analysis. * *p* < 0.05, Friedman’s ANOVA with Dunn’s post-hoc tests (Table S1 for details on statistics). **E**. Classification of poorly and well-trained mice based on their discrimination between go and no-go trials: the further above the 45° line, the better the performance. For this analysis, we used a medial split. **F**. Performance (see STAR Methods) was negatively correlated with the amplitude of the first complex spike peak (r = −0.48, *p* = 0.001, *n* = 45, Spearman rank correlation test). **G**. Also simple spike correlated during the first half of the no-lick period (95-145 ms after trial start) was negatively correlated with performance (r = - 0.40, *p* = 0.006, *n* = 45, Spearman rank correlation test). See also Figs. S1 and S2.

Next, we compared the timing of complex spike firing of naïve mice with that of mice that had completed the training. Given that not all mice reached the same level of choice performance, we took a medial split, resulting in groups with poor and good performance after training (Fig. 3E). The differences in behavior between these groups were particularly evident during the no-go trials: poorly trained mice licked quite often during these trials, more often than the well-trained mice (Fig. 3A-B). The complex spike firing differed between the poorly and well-trained mice: regarding only the go trials, it was evident that the ratio of the complex spike responses to the sound cue and the tactile stimulus was opposite for the poorly and well-trained mice. The poorly trained mice resembled more the pattern of the naïve mice, while the well-trained mice showed a reduced first and an increased second peak in complex spike firing (Fig. 3C; for statistics see Table S1). A correlation analysis revealed that these differential complex spike patterns were related to choice performance, also in the absence of the arbitrary medial split in poor and good performers (r = - 0.48, *p* = 0.001, *n* = 45, Spearman rank correlation; Fig. 3F). A further analysis of all recorded Purkinje cells, irrespective of the relative amplitude of the complex spike responses, yielded similar results (Fig. S1A-B). Thus, the timing of complex spike activity during the trials correlated to choice performance. Complex spike activities during the first window of opportunity, i.e., in response to the sound cue, and those during the second window of opportunity, i.e., related to the tactile cue, were distributed over the network: some Purkinje cells participated mainly during the first peak, others during the second, but there were also Purkinje cells participating in both (Fig. S2A-B). Complex modulation due to the first and/or the second peak was mainly found in the medial part of crus 1 (Fig. S2C).

The simple spike response patterns differed from those of the complex spikes. In naïve mice, the sound cue triggered a double peaked simple spike response with the second peak being more prominent and occurring directly after the initial complex spike response (Figs. 3E and S1C-D). There were no significant differences in simple spike patterns between go and no-go trials in naïve mice (for statistical analysis, see Table S1). In trained mice, simple spike modulation directly following the sound cue was largely absent, irrespective of the task performance and in contrast to the response in naïve mice. The simple spike rate in trained mice with poor choice performance started to increase approximately 100 ms after the sound cue started, whereas in mice with a good performance this moment started later (Fig. 3D and S1C-D). The maximal simple spike modulation during 95-145 ms after trial start – when naïve mice had a clear peak in their simple spike firing – was inversely correlated with behavioral performance (r = −0.40, *p* = 0.006, *n* = 45, Spearman rank correlation; Fig. 3G). During the response window of go-trials, the simple spike rate was increased in trained vs. naïve mice (Figs. 3E, S1C-D), and was particularly obvious in crus 2 (Fig. S2C). Thus, during training, the firing pattern of complex spike increases and subsequent simple spike increases changed together over time: from an early announcing stimulus (i.e., the sound cue) to another stimulus that was initially perceived as neutral and became salient for making a choice (i.e., the rod cue). Accordingly, the changes of both complex spikes and simple spikes correlated to choice performance and in both cases firing frequencies increased towards the period when the mice had to be ready to act and make the proper choice.

### Purkinje cell activity patterns during trials with licking

The previous analyses revealed modulations of complex spike and simple spike firing in relation to the ability of mice to discriminate between go and no-go trials. To study to what extent patterns of Purkinje cell firing could be related to motor execution in trained animals, we subsequently singled out trials with vs. without licks during the response. As we focused here on the licking behavior, we did – for this analysis – not discriminate between go and no-go trials, implying that there were correct and incorrect trials included in this analysis. Sound-triggered complex spikes did occur (infrequently), but they did not differ significantly between trials with and without licks (*p* = 0.784, W = −37, *n* = 34, Wilcoxon matched-pairs test). In contrast, touch-induced complex spikes, occurring later on, did predict whether the mouse would lick afterwards (*p* < 0.001, W = 437, *n* = 34, Wilcoxon matched pairs test; Fig. 4). However, the complex spike activity that predicted the upcoming licking event did not remain elevated during the licking epoch. Simple spike modulation, on the other hand, did not systematically vary before the start of the response window, but was clearly upregulated during the period with licking (Fig. 4, Table S1). Thus, the sequence pattern of complex spike increases and subsequent simple spike increases align temporally with the preparation and execution of the licking behavior, respectively.

**Figure 4.**
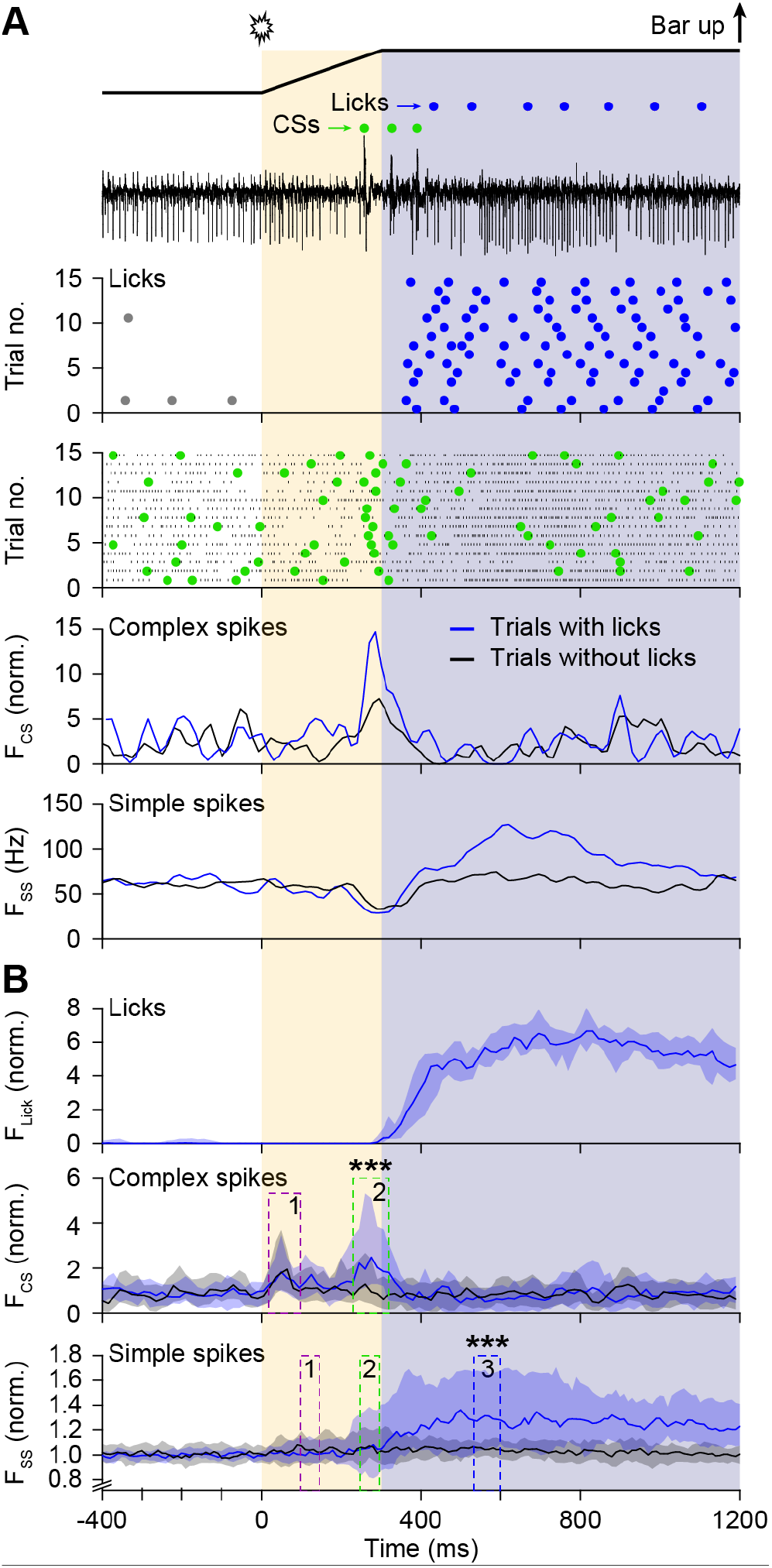
Licking is correlated with complex spike and simple spike activity. **A**. Electrophysiological trace and scatter plots of licks (blue dots), complex spikes (green dots) and simple spikes (black dots) of trials with licking. Below the raster plots are the peri-stimulus time histograms during trials with (blue) and without (black) licking for complex spikes and simple spikes. For this analysis, hit and false alarm trials were grouped together, as were the miss and correct rejection trials. **B**. Peri-stimulus time histograms of 16 mice and 42 Purkinje cells. For complex spike modulation, only Purkinje cells with statistically significant modulation were included (34 cells). The complex spike modulation during the first (sound-evoked) peak was not different between trials with and without licking (*p* = 0.784, W = −37, Wilcoxon matched pairs test), but it was during the second (touch-induced) peak (*p* < 0.001, W = 437, Wilcoxon matched pairs test). Simple spike modulation was not significantly different during the no-lick period, but was during the response window (values for the three intervals indicated in the graph: *p* = 0.812, W = −39; *p* = 0.216, W = 199; *p* < 0.000, W = 577, Wilcoxon matched-pairs test). Lines indicate median values and shaded areas the inter-quartile ranges.

### Complex spike timing is most prominently linked to sensory input

With complex spike activity just before the response window being a good predictor of future licking (Fig. 4), the question arises to what extent the complex spikes reflect the preceding sensory signals and/or the subsequent motor signal that triggers licking. To examine this, we plotted the precise timing of complex spikes relative to the sound cue and licking, respectively, during all hit trials (Fig. 5). In this analysis, we included all recorded Purkinje cells, also those that showed limited modulation in their complex spike rate. Alignment on the sound cue revealed, as expected, a strong resemblance to Figure 4B, with complex spike peaks after the trial onset (i.e., start of the sound cue) as well as just before the start of the response window (Fig. 5A). However, when we aligned complex spike timing to the first lick within the response window (Fig. 5B), we found that the timing of complex spikes was less strict. Thus, the complex spikes were more sharply tuned to the sensory events than to the onset of licking (kurtosis over 500 ms interval: *p* = 0.002, W = 322, *n* = 31 Purkinje cells with statistically significant complex spike peaks, Wilcoxon matched pairs test). In fact, the first lick was even associated with a significant trough in complex spike activity (average complex spike rate −15 to 25 ms around first lick compared to baseline (−700 to −500 ms): *p* = 0.004, W = 292, Wilcoxon matched pairs test). In other words, the precise timing of complex spikes was more tightly coupled to the sensory input than to the motor output.

**Figure 5.**
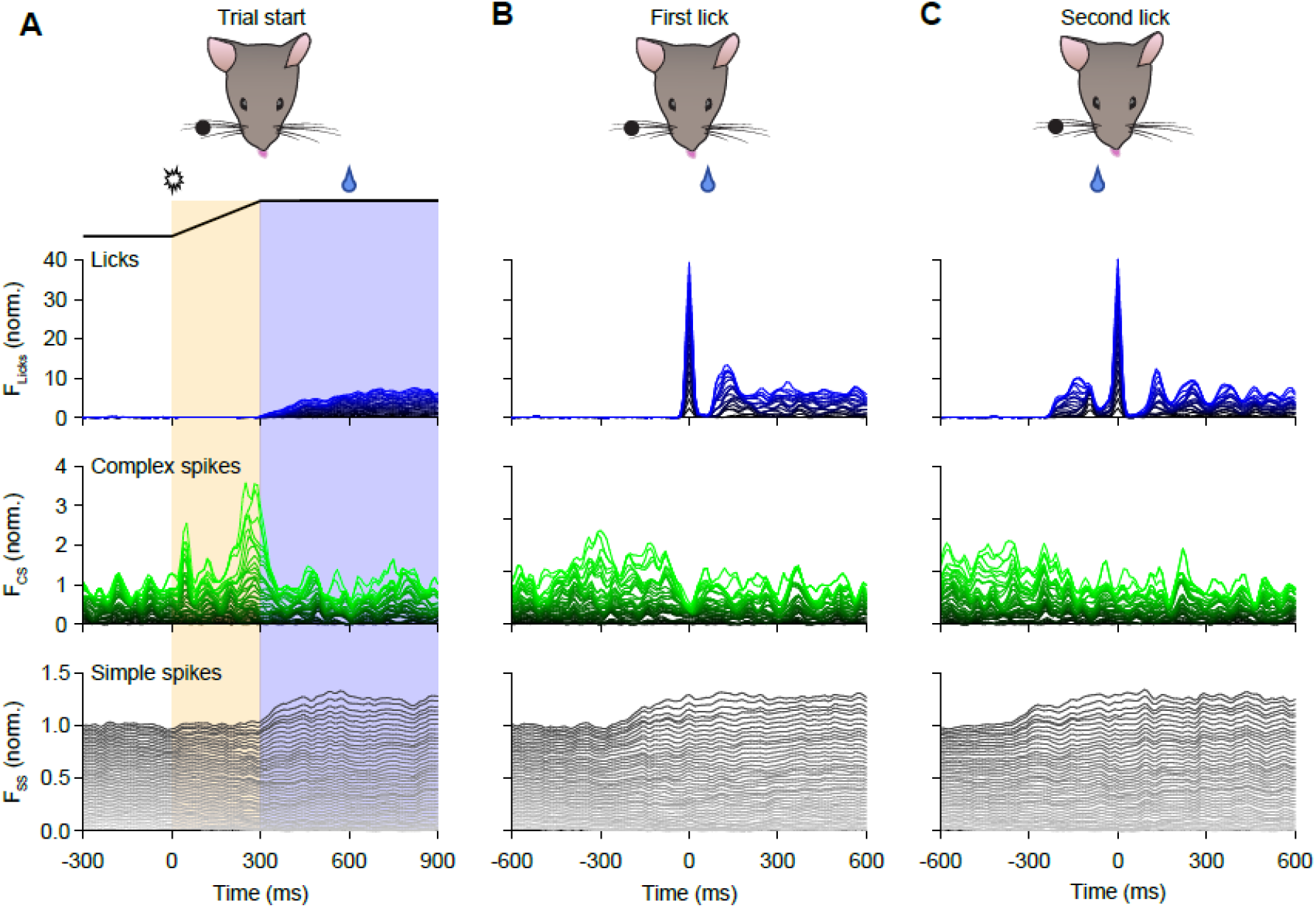
Complex spike timing is mainly linked to sensory input, simple spike timing to motor output. **A**. Stacked line plots of licks, complex spikes and simple spikes during hit trials aligned on the trial onset. The experiments are sorted from smallest to largest modulation and scaled so that the brightest lines indicate the population average. The two complex spike peaks are clearly visible, as is the increase in simple spike firing after the onset of the response window. Data are from 16 mice and 42 Purkinje cells. For this analysis, we did not select Purkinje cells based on their modulation, but included all recorded neurons. **B**. The same data, but now with each trial aligned on the first lick. From this representation it is clear that there is no fixed latency between complex spike firing and the onset of licking bouts. In contrast, the level of simple spike modulation is unaffected and the simple spike increase precedes the licking bout. **C**. The same, but now triggered on the second lick of each trial. As mice had to lick first to obtain a water reward, the mice could expect a water reward during the second lick. The color code and sorting of panels **B** and **C** are based upon the ordering in panel **A**. See also Figs. S3 and S4.

The mice had to lick first before they received a reward. The first lick was therefore unrewarded and, given the potential importance of complex spikes for reward expectation (Heffley and Hull, 2019; Heffley et al., 2018; Kostadinov et al., 2019; Tsutsumi et al., 2019), we repeated this analysis while aligning the complex spikes with the second lick, thus the timing of the reward. This did not reveal a clear coupling between the timing of complex spike firing and reward delivery (Fig. 5C).

To further analyze the impact of complex spike firing just prior to the start of licking, we compared the trials in which a complex spike occurred during the second window of opportunity with those that lacked a complex spike in that time window (240-320 ms after trial start, see Fig. 3C), we did not observe a difference in licking behavior between these trials (Fig. S3). Together, these findings confirmed that the Purkinje cells in crus 1 and crus 2 are tuned to the sensory inputs and trial structure, rather than the actual motor behavior involved in obtaining a reward.

In contrast, the simple spikes showed – in trained mice – predominantly modulation related to the licking behavior, more than to the sensory cues (Fig. 5A-C). It was apparent, however, that there were more simple spikes directly following the second lick – thus at the moment when the mouse noticed that it received a reward – during rewarded than during unrewarded lick bouts (*p* = 0.007, W = 129, *n =* 19 Purkinje cells, Wilcoxon matched-pairs test; Fig. S4). This result is in accordance with previous findings, showing reward-related input to granular cells (Wagner et al., 2017) that can be turned into signals of success at the level of Purkinje cell simple spike encoding (Sendhilnathan et al., 2020).

### Calcium imaging reveals bidirectional evolution of complex spike responses

Our analyses revealed that changes in complex spikes are correlated with performing the timed object detection and choice task. Given their presumptive role in regulating cerebellar plasticity (Coesmans et al., 2004; Gao et al., 2012; Ohtsuki et al., 2009; Romano et al., 2018; Yang and Lisberger, 2014), we assumed that the climbing fiber signals could have a guiding function in cerebellar plasticity of individual Purkinje cells during the learning of our decision task. We therefore wanted to investigate the changes in complex spike activity of individual cells throughout the entire learning process of tens of days. To this end, we repeated the learning experiment in mice in which Purkinje cells were transduced with the genetic Ca^2+^ indicator GCaMP6f in crus 1 using a recently introduced open-source miniscope (de Groot et al., 2020) (Fig. 6A-D).

**Figure 6.**
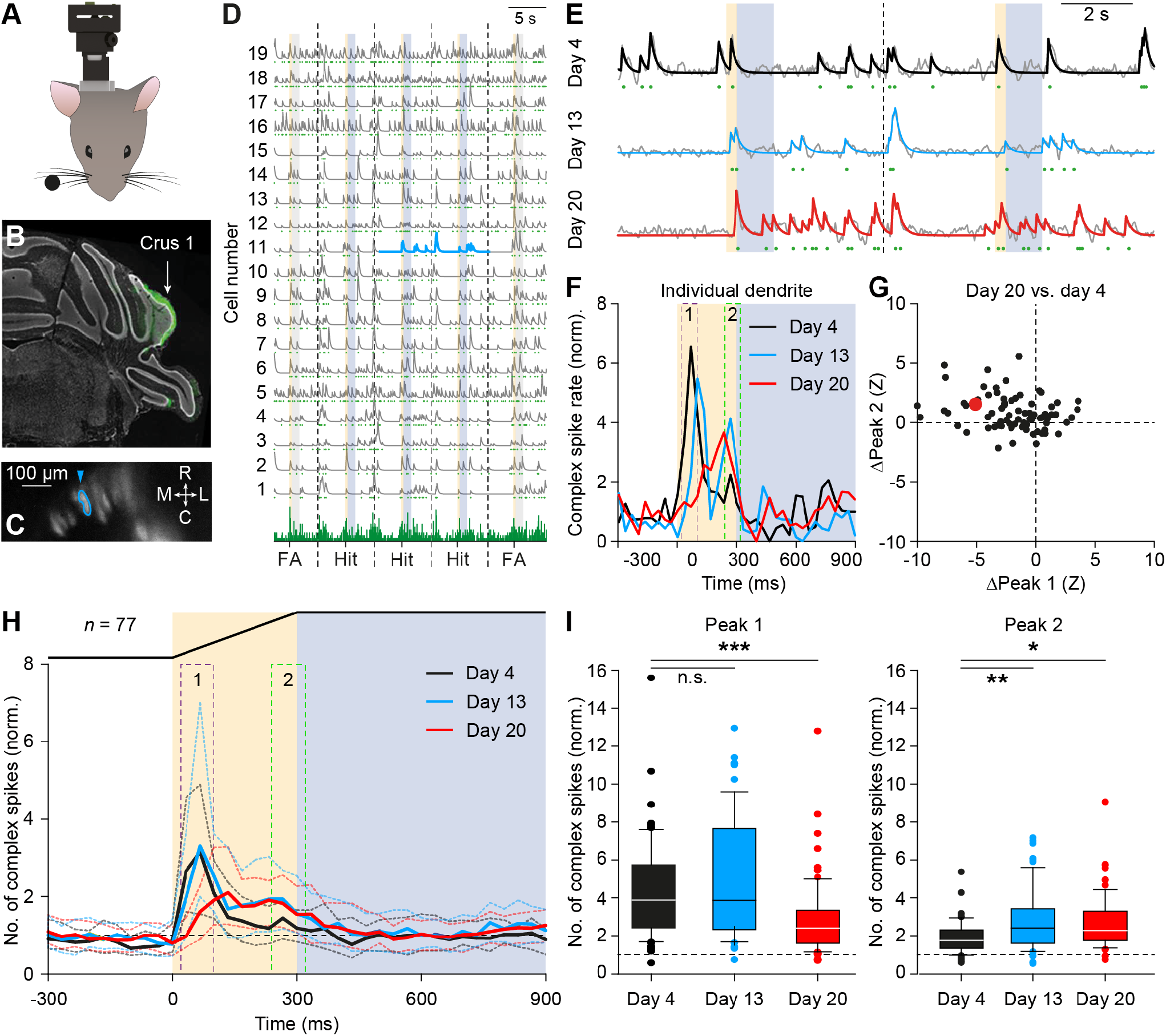
Complex spike plasticity occurs asynchronously. **A**. In four mice, we performed calcium imaging using a miniscope to monitor Purkinje cell calcium transients over the course of training. Purkinje cells were selectively transduced with the genetic calcium indicator GCaMP6f (see STAR Methods). **B**. *Post mortem* histological analysis confirmed the location of transgene expression (using GFP as reporter) in crus 1. **C**. Field of view with 19 dendrites of a representative mouse. The dendrite marked in cyan is highlighted in **D**-**G. D**. Representative recording of 19 dendrites on day 13 of training. The bottom row shows the number of dendrites active at any frame. The light blue fragment is enlarged in **E**. FA = false alarm. **E**. Fluorescent transients of an individual dendrite at days 4, 13 an 20. The grey lines represent the unfiltered trace and the colored lines the convoluted traces. The green symbols in **D** and **E** indicate identified fluorescent transients caused by complex spike firing. **F**. Histograms of the complex spikes of the dendrite illustrated in **E**. Note that the first (sound-evoked) peak changed only during the second half of the training, while the second (touch-induced) peak emerged during the first half. **G**. Scatter plot of the changes in the amplitudes of the first (sound-evoked) and second (touch-induced) complex spike peaks in 77 dendrites that could be identified throughout training. Plotted are the differences between day 20 and day 4. The example dendrite is indicated in cyan. **H**. Median histograms of all 77 dendrites from 4 mice that could be followed throughout training. Dotted lines indicate interquartile ranges. **I**. Box plots showing the complex spike peaks during the first (15-115 ms) and the second (215-315 ms) peaks. First peak: Friedman’s two-way ANOVA (*p* < 0.001, χ^2^ = 20.597, *n* = 77, df = 2) with Dunn’s post-hoc test (days 4-13: *p* = 0.420, χ^2^ = −0.806; days 4-20: *p* < 0.001, χ^2^ = 3.465). Second peak: Friedman’s two-way ANOVA (*p* = 0.003, χ^2^ = 11.403, *n* = 77, df = 2) with Dunn’s post-hoc test (days 4-13: *p* = 0.001, χ^2^ = −3.304; days 4-20: *p* = 0.024, χ^2^ = −2.256; both significant after Bonferroni correction). See also Fig. S5.

In our calcium imaging experiments, we also observed the shift from early (sound-driven) to late (touch-induced) complex spike firing. This was not only visible at the population level, but also in the activity of individual Purkinje cells that could be followed throughout training (Fig. 6E-G). In total, we were able to follow 77 individual Purkinje cells throughout the twenty-day training period. In 36 (47%) of these, both changes occurred, indicating that the double and bidirectional shifts in complex spike activity occur prominently at the level of single Purkinje cells. The decrease of the first (sound-evoked) and the increase of the second (touch-induced) complex spike peaks were asynchronous processes, as already suggested by the exemplary recording in Fig. 6E-F: after 13 days of training, the first peak was not noticeably smaller than a week before (*p* = 0.420, χ^2^ = −0.806, Dunn’s post-hoc test after Friedman’s two-way ANOVA), whereas the second peak had already increased significantly (*p* = 0.001, χ^2^ = −3.304). In the subsequent week, the first peak decreased (*p* < 0.001, χ^2^ = 3.465; Fig. 6H-I). Hence, the learning of the timed object detection task occurred largely in a stepwise fashion. Mice first became more engaged in the task by licking more frequently, but not necessarily at the correct moments. Later in the training, mice began to follow the trial structure with appropriately timed licking (Fig. 2B-C). This seemed to parallel the change in complex spike timing: the recognition of the touch as a salient stimulus occurred before the neglect of the sound.

In our analyses thus far, we ignored the impact of the aversive air puff to the nose that the mice received when they licked during the no-lick interval (see STAR Methods). The reason for this neglect is that the mice performed relatively well during electrophysiological recordings, leaving us with too few trials to reliably analyze. The miniscope recordings enabled us, however, to follow the impact of the aversive puff on complex spike firing. As could be expected, also the aversive puff triggered complex spike firing. Singling out trials with early licks – thus the trials during which the aversive puff was applied – we noticed that also the response to the aversive puff strongly declined with training (Fig. S5A). Even when we used the aversive puff as a trigger, we found hardly any response at the end of the training (Fig. S5B). Thus, the saliency of the aversive puff for evoking complex spike firing was reduced during training, just as for the sound cue.

### Enhancing Purkinje cell simple spike firing during no-response period delays onset of licking

Previous research has shown that a correlation of Purkinje cell activity in crus 1 and crus 2 with licking exists during spontaneous behavior (Bryant et al., 2010). As we showed that a decrease in simple spike firing during the no-response window was associated with choice performance, we wondered to what extent simple spike firing in this period was also directly correlated with licking. To this end, we stimulated Purkinje cells optogenetically in *Pcp2-ChR2* mice centrally in crus 1 and crus 2, thereby creating a temporal disruption in firing of the downstream cerebellar nucleus neurons (Fig. 7A,E) (Romano et al., 2018; Romano et al., 2020; Witter et al., 2013). We did so in mice after training and randomly intermingled trials with and without optogenetic stimulation. When we applied optogenetic stimulation during the no-response period, we noticed that the starts of the licking bouts were delayed in trials with optogenetic stimulation (reduction in number of licks: *p* = 0.007, *t* = 3.503, df = 9, paired *t* test; Fig. 7A-D). This stimulation with a duration of 250 ms had only mild effects on the discrimination between go and no-go trials (fractions of hit and false alarm trials: *p* = 0.055, W = −28; *p* = 0.383, W = 14; Wilcoxon matched pairs tests without correction for multiple comparisons), possibly because the mice had still sufficient time to feel the bar during the 1 s response window that followed after the stimulation period. When we stimulated halfway through the response window, licking was unaffected (*p* = 0.887, *t* = 0.146, df = 9, paired *t* test; Fig. 7E-H). In other words, Purkinje cell activation during the no-response period was related to the start of a licking bout and thereby expression of choice, but when licking was already initiated further enhancing Purkinje cell activity did not have any obvious impact.

**Figure 7.**
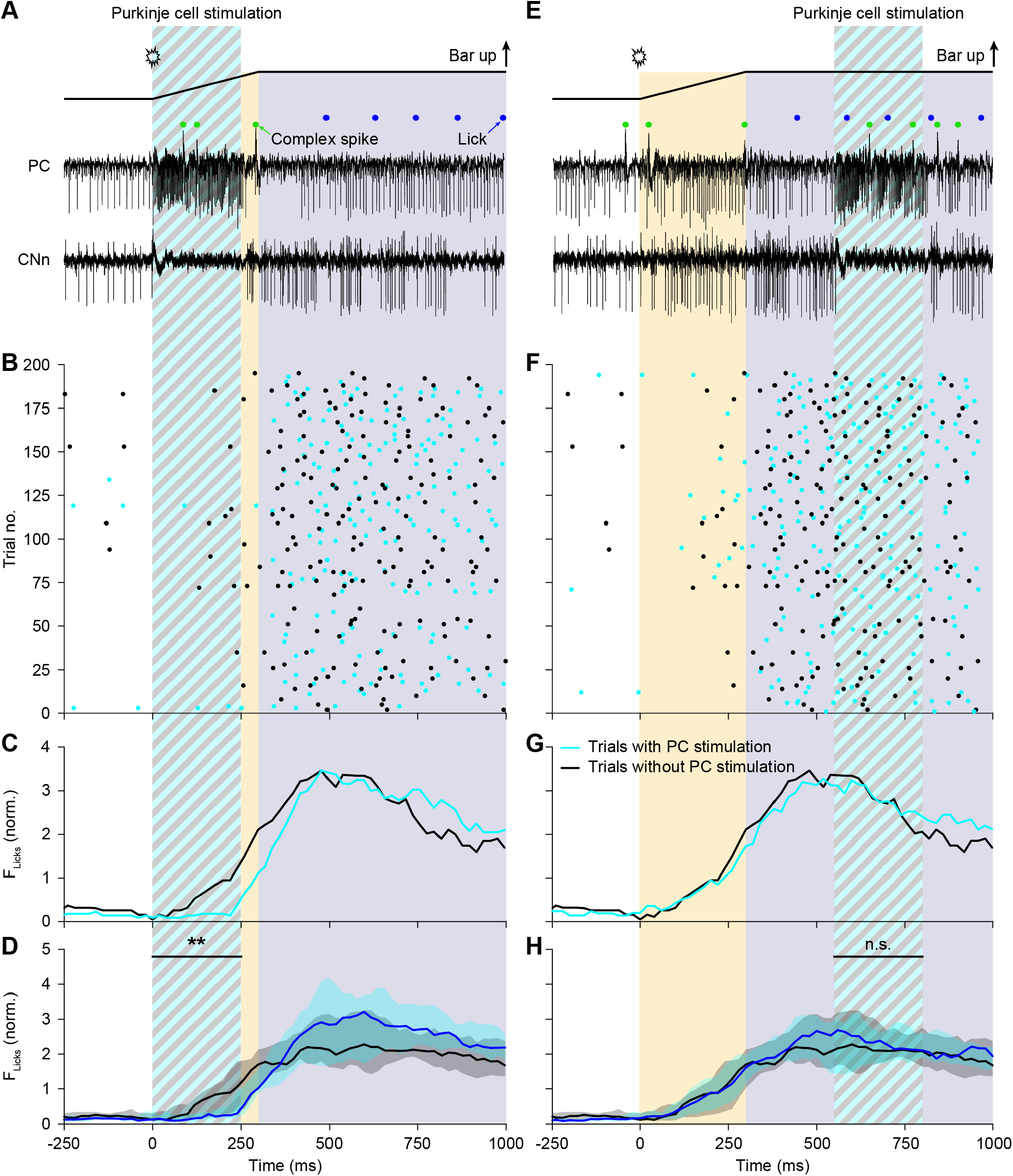
Optogenetic stimulation of Purkinje cells delays the onset of licking, but does not interrupt ongoing bouts. Optogenetic stimulation of Purkinje cells around the border between crus 1 and crus 2 in the lateral hemispheres induced a transient increase in Purkinje cell (PC) simple spike firing and subsequently a decrease in activity of the downstream cerebellar nucleus neurons (CNn). We segregated between stimuli given at the start of licking bouts (**A-D**) and during ongoing licking bouts (**E-H**). Increased simple spike firing induced a delay in the onset of licking, but did not affect ongoing lick bouts. Traces (**A** and **E**) and raster plots (**B** and **F**) are all from the same experiment. In the raster plots, black dots indicate licks during trials without optogenetic stimulation, and cyan dots licks during trials with stimulation. Traces with and without optogenetic stimulation were randomly intermingled. **C** and **E**. Peri-stimulus histograms of the exemplary mouse, and **D** and **H** represent convolved medians of 9 mice with the shaded areas indicating the interquartile ranges.

### Blocking entraining of simple spike increases reduces learning efficacy of choice performance

Given that the pattern of Purkinje cell activity, including both complex spike and simple spike increases, shifts over time during learning of the timed object detection task, our data suggest that the changing activity patterns of Purkinje cells in crus 1 and crus 2 are tightly related to the structure of the trial, guiding the choices. Given that the increases in simple spike activity appear to consistently follow those of the complex spikes, one wonders to what extent the impact of both acquired changes, i.e., that of the complex spikes and that of the simple spikes, can be disentangled. We thereto studied the learning behavior as well as Purkinje cell activity during the timed object detection task in *Pcp2-Pppr3r1* KO mice (Fig. 8A), which have been reported to suffer from changes in simple spike activity due to a deficiency in postsynaptic potentiation at the parallel fiber to Purkinje cell inputs, while leaving the complex spike activity intact (Schonewille et al., 2010). The *Pcp2-Pppr3r1* KO mice required significantly more time (fraction correct trials: *p* = 0.006, *F* = 3.215, df = 17, interaction effect, repeated measures ANOVA with Greenhouse-Geisser correction) to master the discrimination task than wild type mice (Fig. 8B). In contrast, the *Pcp2-Pppr3r1* KO mice were still able to execute the licking movements at a normal performance level (Fig. 8C). The complex spike activity of Purkinje cells in crus 1 and 2 of *Pcp2-Pppr3r1* KO mice appeared unaffected during spontaneous behavior, whereas the simple spike activity was reduced and more regular (Fig. S6A), the latter in agreement with previous findings (Rahmati et al., 2014; Romano et al., 2018; Schonewille et al., 2010). Likewise, when we analysed the complex spike activity during the hit trials of the timed object detection task (Fig. 8C), we did not observe significant differences between wild types and mutants (Fig. 8D). The timing of the complex spike activity was not much affected, with neither the amplitude of the first, nor that of the second peak being significantly different between wild type and *Pcp2-Pppr3r1* KO mice (*p* = 0.462, U = 58.5 and *p* = 0.388, U = 56, respectively, Mann-Whitney tests). In contrast, the simple spike pattern was completely different in *Pcp2-Pppr3r1* KO mice. Rather than a broad increase in simple spike firing during the response window as occurs in wild types, *Pcp2-Pppr3r1* KO mice showed a decrease in simple spike firing (e.g. during 545-605 ms: *p* < 0.001, U = 58, Mann-Whitney test; Fig. 8E). Thus, the mutant mice that are deficient in Purkinje cell long-term potentiation (Romano et al., 2018; Schonewille et al., 2010) indeed fail to show an upregulation of simple spike firing during learning. This effect appeared to be specific for the response stage, because the simple spike firing during the early (non-response) phase of the trial was equally low in trained wild type and *Pcp2-Pppr3r1* KO mice (95-145 ms: *p* = 0.826, U = 18.5, Mann-Whitney test; Fig. 8E). These data suggest that the changes in simple spike activity that follow the changes in complex spike activity also contribute to choice performance.

**Figure 8.**
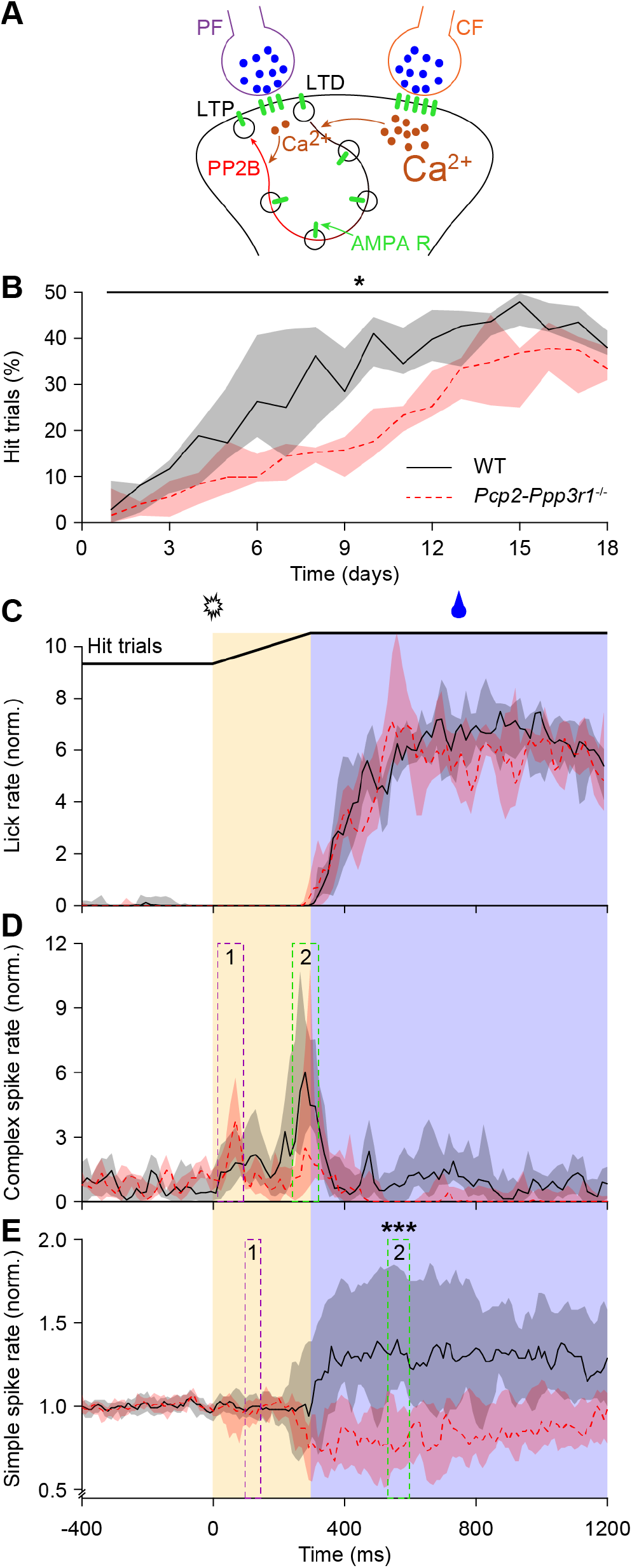
Impairment of Purkinje cell LTP reduced learning efficacy, but not motor performance in trained mice. **A**. *Pcp2-Ppp3r1*^-/-^ mice lack protein phosphatase 2B (PP2B) specifically in their Purkinje cells. As a consequence, the Purkinje cells of the mutant mice are not able to express parallel fiber-to-Purkinje cell LTP. Co-activation of parallel fibers (PF) and the climbing fiber (CF) leads to a large influx of Ca^2+^ into Purkinje cells, favoring parallel fiber-to-Purkinje cell LTD, while activation of parallel fibers in the absence of climbing fiber activity induces LTP, involving activation of PP2B in wild type mice. Schematic drawing adapted with permission from (Romano et al., 2018). **B**. During training, *Pcp2-Ppp3r1*^-/-^ mice took a longer time to learn to time their licks than their wild type controls. Medians (shades: interquartile ranges) of the fraction of hit trials in 9 mutant mice and 9 wild type littermates. Peri-stimulus histograms (medians with interquartile ranges) of the number of licks (**C**), complex spikes (**D**) and simple spikes (**E**) during hit trials of WT and mutant mice. The complex spikes were taken only from significantly responsive cells (9 in mutant mice and 16 in wild type mice), the simple spike from all 14 and 27 cells, respectively. Although the actual motor performance was comparable, the simple spike modulation was different (95-145 ms: *p* = 0.826, W = 180.5; 545-605 ms: *p* < 0.001, U = 58, Mann-Whitney tests of mutant vs. wild type cells). See also Fig. S6.

## Discussion

How do we learn to make targeted meaningful movements the way we do, combining cognitive with motor functionalities over time? To optimize movements during complex tasks, our brain continuously makes predictions on the outcomes of our actions. The olivocerebellar system plays a critical role in creating such expectations and adapting movements on the basis of sensory feedback (Brooks et al., 2015; Cayco-Gajic and Silver, 2019; Heffley et al., 2018; Hull, 2020; Kostadinov et al., 2019; Larry et al., 2019; Moberget and Ivry, 2019; Tsutsumi et al., 2019; Tzvi et al., 2020; Wolpert et al., 1998). A specific and particularly well-studied example of a relatively simple form of cerebellar motor learning is eyeblink conditioning: subjects can learn to associate an initially neutral stimulus, such as a sound or an LED light, with an air puff to the cornea, and eventually close their eyelids upon perceiving only the initially neutral stimulus (Koekkoek et al., 2003; Ohmae and Medina, 2015; Steinmetz et al., 1987; Ten Brinke et al., 2015). Conversely, as done in the current study, more complex, go/no-go detection study, subjects can also learn to associate an initially neutral stimulus with a pleasant outcome, such as the delivery of a water drop. In either case, the climbing fibers of the olivocerebellar system apparently shift the identity and timing of their signaling in that they increase their responses to the initially neutral sensory stimulus, triggering the conditioned movement, and that they start to arise at the moment when this movement emerges, engaging a readiness to act. During both the simple eyeblink conditioning paradigm and the complex go/no-go detection task, the novel complex spike response rises while their responses to the unconditioned stimulus reduce. This shift in saliency and timing can occur independently from the sequence of events in that the initially neutral stimulus can occur either before (as in eyeblink conditioning) or after (as in the go/no-go task) the unconditioned stimulus. Moreover, the shift in saliency and timing of the acquired complex spike response can also occur independently from the polarity of the associated subsequent simple spike response in that this can either decrease (as in eyeblink conditioning) or increase (as in the go/no-go task) directly after the emerging complex spike response. The shift in both the complex spike and simple spike responses over time correlates well with the choice performance of the go/no-go detection task, but at the end of the training the complex spike response reflects more the saliency of the sensory stimuli and a readiness to act, whereas the simple spike response is closer related to the ongoing motor response. Our chronic calcium-imaging recordings of multiple Purkinje cells across tens of days revealed that the shift in saliency and timing of the complex spike activity during the go/no-go detection task occurs prominently at the level of individual cells, yet they also showed that the shift takes time and comprises a stage during which the climbing fibers of individual cells robustly respond to both the unconditioned stimulus and the initially neutral stimulus. Even though such chronic imaging recordings have not been made during eyeblink conditioning, acute electrophysiological recordings suggest that the same phenomena may occur over time during this paradigm (Ohmae and Medina, 2015; Ten Brinke et al., 2019; Ten Brinke et al., 2015). Thus, the universal mechanism during both forms of learning may be that the climbing fiber signals move to the most salient sensory input that engages a readiness to act, which is expressed by a subsequent alteration in simple spike activity. As a consequence of this mechanism, the olivocerebellar system appears to be able to coordinate the non-motor component of a complex task over time to its motor component.

### Go/no-go detection task for testing decision-making

During the go/no-go task that we designed the mice had to learn to make a decision to act or not upon perceiving a tactile cue. In doing so, the mice had to learn to ignore the auditory cue that was not predictive of the presence or absence of a reward. Our timed object detection task differed from other go/no-go paradigms (Rahmati et al., 2014) in that we included a no-lick period of 300 ms between trial onset and the start of the response window. Thereby our paradigm allowed us to precisely relate Purkinje cell activity to the two sensory cues as well as to the actual motor action. Our paradigm also differed from other studies that investigated the cerebellar role in reward expectation in that those studies essentially trained associative learning between a stimulus or action and the direct presence of a reward (Heffley et al., 2018; Kostadinov et al., 2019; Larry et al., 2019; Tsutsumi et al., 2019; Wagner et al., 2017). In our paradigm, mice were trained to respond to a tactile stimulus and received a reward only when they reacted at the right time during go trials. Importantly, the reward was not instantly present, but only administered after the mice licked first. The presentation of a reward could therefore at least not directly guide the behavior. Thus, our paradigm required decision-making to engage an action, highlighting a global role for the olivocerebellar system that extends beyond reward expectation learning.

### Preparatory motor activity in the cerebellum

Motor planning and expectation are closely related, so it is natural to assume that the cerebellum participates in preparatory motor activity. Indeed, the cerebellum acts in a loop with the anterolateral motor cortex during motor planning (Chabrol et al., 2019; Gao et al., 2018). Such preparatory motor activity is typically expressed by ramping or sustained increased activity of cerebellar nuclei neurons (Chabrol et al., 2019; Gao et al., 2018). As Purkinje cells provide strong inhibition to the neurons of the cerebellar nuclei (Ito et al., 1964), it comes as no surprise that many of them decrease their simple spike firing during preparatory motor activity (Chabrol et al., 2019). In a minority of cells, the opposite pattern has been described: increased simple spike firing by Purkinje cells and inhibition of cerebellar nucleus neurons (Chabrol et al., 2019; Gao et al., 2018). A generalization of the interpretation of preparatory activity in the cerebellum is not straightforward though, as under many conditions preparatory activity can be confused with sensory responses or with ongoing complementary behavior (De Zeeuw and Ten Brinke, 2015). In our wild type dataset, we found that the shift in timing of the simple spike response co-occurred with that of the complex spike response. Once the complex spikes respond to the more salient stimulus, the simple spike increase that directly followed moved concomitantly with that of the complex spikes, highlighting the preservation of a particular complex spike – simple spike sequence, but shifted over time during acquisition of the detection task. Indeed, as learning progressed the initial simple spike peak during the 300 ms delay (non-response) period was minimized, while the secondary simple spike response during the response period emerged, which was persistent during the actual licking response, suggesting an ongoing contribution.

### Motor activity in the cerebellum

To what extent do the Purkinje cells in crus 1 and crus 2 have a motor function in the timed object detection task? During spontaneous licking, thus in the absence of a training or timing context, simple spikes in these lobules modulate with rhythmic licking (Bryant et al., 2010). Simple spikes have different phase relations with licking, reminiscent of what occurs during other rhythmic processes such as walking (Sauerbrei et al., 2015) or breathing (Romano et al., 2020). Transient or permanent lesions of the cerebellum result in licking with a lower frequency, suggesting a functional contribution of Purkinje cells to generating motor output (Bryant et al., 2010). Yet, considering the cerebellum as an internal model for generating prediction errors, Purkinje cells may predominantly receive motor efference copies and contribute relatively little to the initiation of unperturbed motor activity (Wolpert et al., 1998), which is in line with the timing of simple spike modulation during unperturbed breathing as well as naïve whisker reflexes (Romano et al., 2018; Romano et al., 2020).

In our current study, several arguments argue against a dominant driving role for Purkinje cells in crus 1 and crus 2 during timed licking in a go/no-go task. First, at the end of the training complex spikes were more sharply tuned to the sensory cues than to the initiation of motor behavior, while the simple spikes occurred predominantly after licking onset. Second, the secondary simple spike response in *Pcp2-Pppr3r1* KO mice that emerges during the training showed a decrease rather than an increase during licking, while the ability to lick was, unlike the choice performance, unaffected. Finally, trial-by-trial analysis of the complex spikes indicated that individual complex spikes did not contribute significantly to the licking behavior, neither in the same trial nor in the subsequent trial. Thus, instead of supporting a direct role in driving the motor activity during our go/no-go detection task, our findings rather point towards an overall contribution of Purkinje cells in crus 1 and crus 2 to a general engagement to act.

### Changes in Purkinje cell activity correlate with learning

During learning the complex spike and simple spike patterns change concomitantly and are correlated with choice performance. *Pcp2-Pppr3r1* KO mice show reduced learning efficacy, suggesting that Purkinje cell potentiation – which is absent in these mice (Schonewille et al., 2010) – is required for efficient learning of our timed object detection task. Their learning deficits are in line with the absence of an entrained increase in their simple spike firing during the response window.

Classical cerebellar theory predicts that the co-occurrence of climbing fiber and parallel fiber activity leads to LTD of the parallel fiber-to-Purkinje cell synapse (Albus, 1971; Ito, 2002), the strength of which depends on the relative timing of climbing fiber and parallel fiber activation as well as the intrinsic properties of the local Purkinje cells involved (Suvrathan et al., 2016). In naïve mice, the sound cue at the start of a trial triggered both complex spikes and simple spikes, which may well have led to the induction of LTD. This assumption is compatible with the finding that the simple spike response to the sound cue was strongly reduced during learning. Instead, later on in the trials, the simple spike rate increased during the response windows during which little or no complex spikes were triggered (albeit they did occur just before this period). This could be expected, as the absence of climbing fiber activity provides a window of opportunity for simple spike potentiation (Coesmans et al., 2004; Romano et al., 2018). Moreover, this possibility is supported by the observation that Purkinje cell-specific PP2B mutants, which lack LTP induction (Rahmati et al., 2014; Romano et al., 2018; Schonewille et al., 2010), show impaired learning of paradigms that are mediated by downstream pathways in which the net polarity of synaptic connections in the network is inhibitory (De Zeeuw, 2020; Romano et al., 2018; Voges et al., 2017).

Calcium imaging and behavioral analysis suggest that the depression and potentiation mechanisms form indeed two complementary processes. Mice first learn to lick during the trials, and then learn to time the licks (Fig. 2B-C). The sound-evoked complex spikes, abundant during the start of the no-lick interval in naïve mice, start to decrease relatively late during training (Fig. 6H-I), and remain profusely present in mice with a poor learning efficacy (middle panel, Fig. 3C). The latter is coupled to an incomplete reduction of simple spike firing during the first half of the no-lick window (Fig. 3D), suggesting an imperfect LTD.

In the meantime, the touch-induced complex spikes, occurring during the last phase of the no-lick period, are already upregulated during an early phase of training (Fig. 8E-F). However, as the sound cue is still perceived as salient (as evidenced by the strong complex spike response), the mice still have trouble with timing. Mice with a slow learning rate, whether wild type or *Pcp2-Pppr3r1* KO mice, did not have problems with licking – they were impaired in learning to lick at the right moment and this deficit occurred only during go trials. These experiments suggest that individual Purkinje cells in the lateral cerebellum can combine complementary plasticity mechanisms induced by the changing presence and absence of their climbing fiber input over time during the training task and thereby shift not only the pattern of complex spike activity, but also that of the simple spikes. As a consequence, these Purkinje cells can set the stage for well-timed, context-dependent behavior, engaging a readiness to act, but without being indispensable for acute motor execution, nor being restricted to reward expectation. Together, in line with the periodic operations of neurons in the inferior olive (Negrello et al., 2019), our findings highlight how climbing fiber signaling in the olivocerebellar system may set the pace when coordinating non-motor with motor functions.

## Methods

### Animals

*Pcp2-Pppr3r1* (“L7-PP2B”) mice (Tg(Pcp2-cre)2MPin;Ppp3r1^tm1Stl^) lacked functional PP2B specifically in their Purkinje cells. They were created by crossing mice in which the gene for the regulatory subunit (CNB1) of PP2B was flanked by loxP sites (Zeng et al., 2001) with transgenic mice expressing Cre-recombinase under control of the *L7* (*Pcp2*) promoter (Barski et al., 2000) as described in Schonewille et al. (2010). Learning curves of L7-PP2B mice (4 males and 5 females) were compared with their control littermates (5 males and 4 females) trained together. Optogenetic experiments as well as Purkinje cells recordings from naïve mice were performed on transgenic mice that expressed channelrodopsin2 (ChR2, 11 males and 3 females) also under the *Pcp2* promoter (Witter et al., 2013). The animals were group housed until magnetic pedestal placement; after that they were single housed in a vivarium with controlled temperature and humidity and a 12/12h light/dark cycle. All recordings and behavioral experiments were performed in awake, head restrained mice with an age between 11 and 35 weeks. All mice were healthy and specific pathogen free (SPF). All experimental procedures were approved *a priori* by an independent animal ethical committee (DEC-Consult, Soest, The Netherlands) as required by Dutch law and conform the relevant institutional regulations of the Erasmus MC and Dutch legislation on animal experimentation. Permission was filed under the license numbers EMC3001, AVD101002015273 and AVD1010020197846.

### Habituation and water restriction

Mice received a magnetic pedestal for head fixation, attached to the skull above bregma using Optibond adhesive (Kerr Corporation) under isoflurane anesthesia (2–4% v/v in O_2_). Postsurgical pain was treated with carprofen (Rimadyl, Pfizer) and lidocaine (Braun) and two days of recovery followed the procedure. In order to reduce and standardize stress level during training, the experimenter began to handle mice a week before the start of the actual training for approximately 15 minutes per mouse per day. Starting from three days before the training, the water bottles were removed from the lid of the cages and the body weight of mice was daily monitored and mice were head fixed and restrained for 15 minutes each day; during this time, water was available from the lick-port positioned in front of the mouse. Mice that did not drink during this time received anyway a controlled amount of water in their cages; in total mice received a daily amount of 1 ml of water per 20 g body weight.

### Behavioural paradigm and surgical procedures

Mice were trained for 18 days to associate the position of a pole with the presence or absence of a water reward. During training, go and no-go trials were randomly intermingled. In go trials, a pole was raised in the middle of the whisker field and the mice received a water reward when they licked during the response interval. An important difference with paradigms generally described in other works (e.g. (Heffley and Hull, 2019; Heffley et al., 2018; Kostadinov et al., 2019; Larry et al., 2019; Tsutsumi et al., 2019)) is that the water delivery was triggered by the first lick of bout, in the moment when the laser beam in front of the lick-port was interrupted; as a result, mice had to lick before the reward was given, so that they could not use the presence of water nor the valve click as a cue. During no-go trials, mice were not supposed to lick and licking was consequently not rewarded. Each trial, whether go or no-go, started with a clearly audible sound made by the pneumatic device raising the pole. The pole was constructed so that the location and the characteristics of the sound were identical between go and no-go trials. Licking during the 300 ms period following trial start, announced by the sound cue, was not allowed and induced early termination of the trial and an aversive air puff to the nose of the mouse. During the first two days of training the aversive puff was omitted to facilitate the participation to the task. During training, body weight and health condition of mice were monitored and mice not cooperating or not in good health condition were taken out of the experiment (5 out of 40).

At the end of the training mice received the water bottle in their cages for two days. Once recovered from the water restriction regime, a craniotomy was performed to expose cerebellar crus 1 and crus 2; being this procedure longer and more invasive than the pedestal placement, the analgesia previously mentioned was complemented with bupivacaine (Actavis, Parsipanny-Troy Hills, NJ, USA) and buprenorphine (“Temgesic”, Indivior, Richmond, VA, USA), and the recovery period was three days long. The craniotomy was cleaned and covered with Kwik-Cast (World Precision Instrument, Sarasota, FL, USA).

After mice recovered, the water restriction regime restarted. A retraining phase of 2 to 5 days preceding the electrophysiology allowed us to verify, apart from the health condition of our mice, that the participation level was suitable for efficient electrophysiological recordings.

### Optogenetic stimulation

After craniotomy and retraining, *Pcp2*-*Cre*/*Ai27* mice underwent to two task sessions (consecutive days) during light stimulation. An optic fibre (diameter 400 µm, Thorlabs, Newton, NJ, USA) was placed in the middle of the craniotomy perpendicular to the cerebellar surface. Three conditions were randomly intermingled for both go or no-go trials: a control condition of unaltered go or no-go trials and two conditions where a pulse of blue LED light (λ = 470 nm, duration = 250 ms, P = 5 mW) was given. During the first session, the light pulse was delivered either at time 0 ms (together with the acoustic cue) or after 300 ms (when the pole reached the top position); during the second session the light turned on either at 0 ms or at 550 ms during the response window, when licking was generally already ongoing. After the session the craniotomy was rinsed with saline and closed with Kwik-Cast. Purkinje cells of these mice were recorded in a subsequent session one to three days later.

### Electrophysiology

Electrophysiological recordings were performed in awake mice using quartz-coated platinum/tungsten electrodes (R = 2-5 MΩ, outer diameter = 80 µm, Thomas Recording, Giessen, Germany). Electrodes were placed in an 8×4 matrix (Thomas Recording), with an inter-electrode distance of 305 µm. Prior to the recordings, the mice were lightly anesthetized with isoflurane to remove the dura mater, bring them in the setup and place the electrodes on the surface of the cerebellum. Recordings started at least 60 min after termination of anaesthesia and were made in crus 1 and crus 2 ipsilateral to the side of the whisker stimulation at a minimal depth of 500 µm. The voltage signal was digitized at 25 kHz, using a 1-6,000 Hz band-pass filter, 22x pre-amplified and stored using a RZ2 multi-channel workstation (Tucker-Davis Technologies, Alachua, FL). Once awake, mice attention was triggered by randomly delivering few drops of water until they spontaneously started seeking for water. Once good stable signal was found from at least one cells and anyway not after more than 90 minutes from the moment we remove the anaesthesia, the behavioural session was started and continued until mice stopped drinking and we collected a certain amount of trials in the absence of licking responses.

### Behavioral data analysis

Licking bouts were defined as sequences of licks with intervals <500 ms. Trials in which the trial start fell within an ongoing licking bout (that started at least 20 ms before the sound cue was given) were ignored for the calculation of performance. Learning performance was calculated as the ratio between hit trials and the sum of false alarm and early lick trials.

After optogenetic stimulation we were interested in observing if any changes were induced by the light in the licks’ distribution following the cues or within ongoing bouts. We therefore built peri-stimulus time histograms (PSTHs, 50 ms bins) of the latencies of licks from trial onset with and without light stimulation, then compared the licking probability in the two 250 ms windows starting from 0 ms or 550 ms.

### Electrophysiological data analysis

Spikes were detected offline using SpikeTrain (Neurasmus, Rotterdam, The Netherlands). A recording was considered to originate from a single Purkinje cell when it contained both complex spikes (identified by stereotypic waveform, overshooting and the presence of spikelets) and simple spikes, and in which each complex spike was followed by a pause of at least 8 ms before simple spike firing resumed. When comparing two or more conditions, only recordings containing at least 8 events per condition were included in each group. We generally used bins of 10 ms to visualize simple spikes and of 15 ms for complex spikes and licks. In order to compare the modulation from different cells or evoked by different triggers, PSTHs have been normalized on the average firing frequency calculated in a 2 seconds interval preceding the second before the trigger. To compare complex spikes modulation to trials cues, peaks have been detected in the two temporal windows of interest (20-100 ms and 240-320 ms after trial start) as maximum bin value. A cell was considered modulating when in one or both the temporal windows the maximal complex spike modulation exceeded at least 3 standard deviations the average baseline frequency. Simple spikes virtually always showed some degree of modulation, so that we did not separate them into responsive and non-responsive cells.

### Miniscope imaging

Calcium transients were imaged daily in a group of four mice using the NINscope miniscope using procedures described previously (de Groot et al., 2020). Briefly, mice were anesthetized with isoflurane in a stereotactic apparatus and a pedestal for head fixation was mounted. A 2 mm round craniotomy was made centred above cerebellar lobule crus 1 to inject virus (AAV1.CAG.FLEX. GCaMP6f/AAV1.CMV.PI.Cre.rBG mixed 1:1, which was diluted 1:3 in saline) for transduction of Purkinje cells with GCaMP6f, and to mount a gradient index (GRIN) lens. Fifteen minutes prior to virus injection, D-mannitol (15% in saline) was injected i.p. to facilitate virus diffusion (Kuhn et al., 2012). Virus was injected at four locations. At each location 25 nl of virus was injected once at 350, twice at 300 and once at 250 µm depth at a rate of 25 nl/min with a Nanoject II Auto-Nanoliter Injector (Drummond Scientific Company, USA). After injection of the virus, a 1.8 mm GRIN lens was implanted. Kwik-Sil (WPI, USA) was applied around the edges of the craniotomy and the lens. Subsequently, the lens was secured by applying dental cement (Super-Bond C&B, Sun Medical, Japan). The lens was covered with Kwik-Cast (WPI, USA) for protection. Two to three weeks after viral injection a baseplate was mounted in an optimal location and secured with dental cement.

Before training commenced, mice were first habituated for a week to being head-fixed using the head pedestal and for the mice to discover the location of the lick-port and water reward. Mice were then subjected to the same training protocol as described before, but now with a mounted miniscope for calcium imaging. For every session, 220 frames were collected at 30 Hz. Recordings began 3 seconds before presentation of the first stimulus. Imaging continued for a period of twenty days from commencement of training.

### Extraction and analysis of calcium transients

Raw data were motion-corrected using noRMCorre (Pnevmatikakis and Giovannucci, 2017) and calcium transients were extracted using CNMF-E (Zhou et al., 2018). In order to compare the modulation of the same cells across three different training sessions motion corrected frames recorded at days 4, 13 and 20 of training were concatenated in order to obtain one large video; between-sessions misalignment was corrected using ImageJ: the frames composing each session were averaged, then the three averages were manually overlapped and the exceeding pixels on the x and y axis were cropped from each frame. We ran CNMFE on these aligned data to extract spatial footprints and signals of Purkinje cell dendrites. Variations in the baseline signal present across different sessions were subtracted (mean of sliding median and sliding minimum, 25 frames sliding window). Deconvolved transients were used to determine the onset of the calcium transients. Bin size for peri-stimulus histograms was set at 0.0332 seconds given a 30 Hz acquisition.

## Quantification and statistical analysis

Statistical tests employed are mentioned throughout the manuscript. When applicable, corrections for multiple comparisons have been applied. This is indicated in the text. Tests were two-sided.

## Acknowledgements

The authors wish to thank Sander van Gurp, Sander Lindeman and Mario Negrello for their contributions to the pilot phase of this project, Hugo Hoedemaker, Maryam El Hamdioui and Danique Broere for their aid with surgeries and data collection, and Sanne Smid for help with video analysis. Financial support was provided by the Netherlands Organization for Scientific Research (NWO-ALW; CIDZ), the Dutch Organization for Medical Sciences (ZonMW; CIDZ), BIG (CIDZ), Medical Neuro-Delta (CIDZ), INTENSE LSH-NWO (CIDZ), ERC-adv and ERC-POC (CIDZ), Van Raamsdonk-fonds (CIDZ), 3V-Fonds KNAW (TH and CIDZ), and Albinism Fonds NIN (CIDZ).

## Author contributions

Conceptualization: L.B., L.W.J.B. and C.I.D.Z.; Methodology: L.B., T.M.H. and L.W.J.B.; Software: L.B., T.M.H. and L.W.J.B.; Validation: L.B., T.M.H. and L.W.J.B.; Formal Analysis: L.B.; Investigation: L.B. and V.R.; Resources: T.M.H. and L.W.J.B; Data Curation: L.B.; Writing – Original Draft: L.B. and L.W.J.B.; Writing – Review & Editing: T.M.H., L.W.J.B. and C.I.D.Z.; Visualization: L.B., T.M.H. and L.W.J.B.; Supervision: L.W.J.B. and C.I.D.Z.; Project Administration: L.B. and L.W.J.B.; Funding Acquisition: T.M.H. and C.I.D.Z.

## Supplemental Information

**Figure S1.**
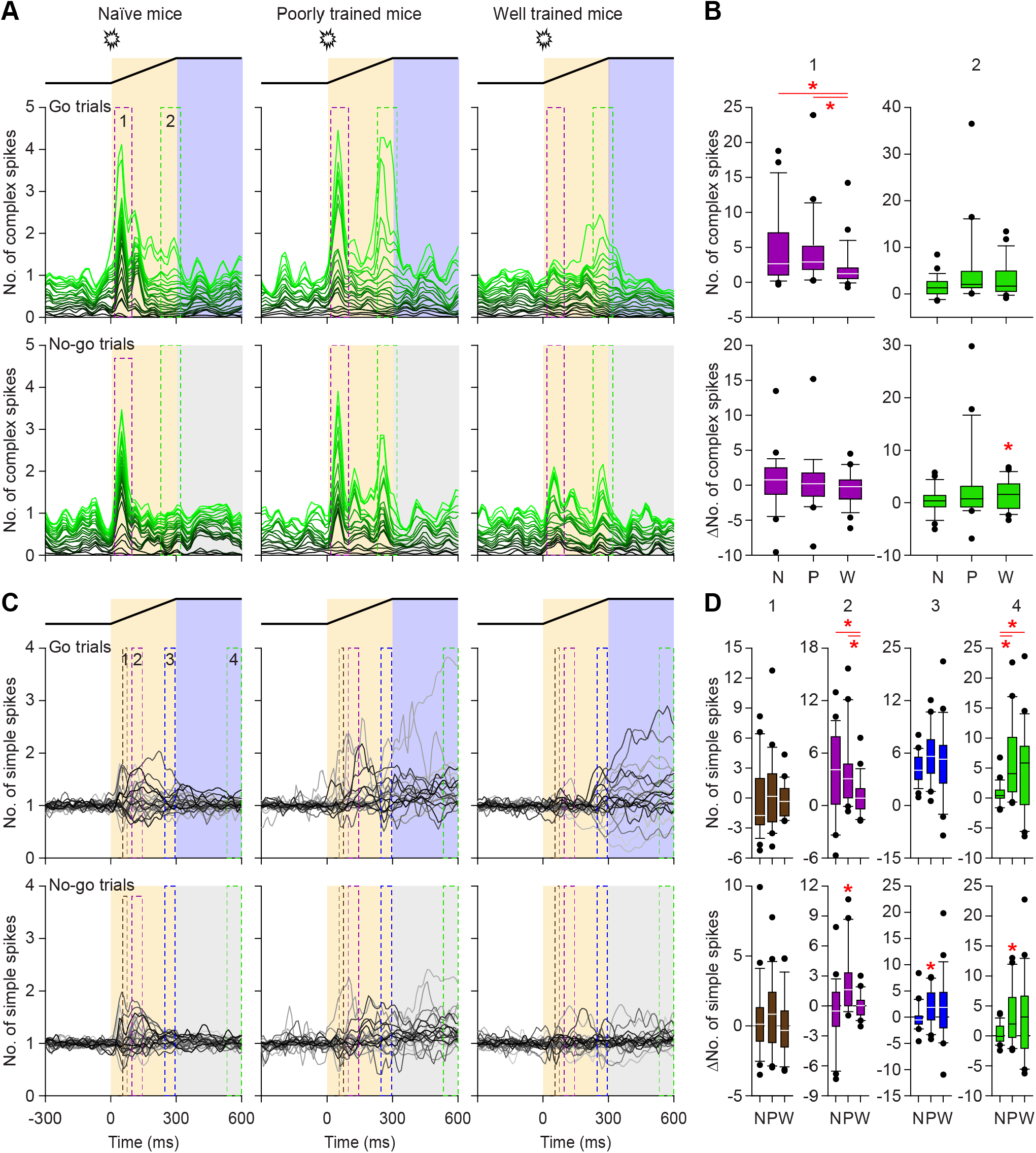
Changes in complex spike and simple spike timing during learning, belonging to Fig. 3. **A**. Stacked line plots of complex spike firing in naïve (left), poorly trained (middle) and well-trained (right) mice (see Fig. 3E) during go and no-go trials. Each line represents the peristimulus time histogram of a single Purkinje cell. The Purkinje cells are sorted based upon the maximal complex spike response during go trials and normalized to the pre-trial activity and scaled so that the upper (brightest) line represents the population average. It is clear that the first (auditory) cue at the start of the trial has a stronger impact than the second (tactile) cue. All 24 recorded Purkinje cells are included in this analysis, irrespective of whether they displayed a statistically significant response. **B**. Top row: box plots of the maximal complex spike peak during the first (20-100 ms, left) and the second (240-320 ms, right) time window. Bottom row: box plots of the difference in maximal response for the for the first (left) and second (right) time window between go and no-go trials. **C** and **D**. The same for the simple spike activity. For simple spikes, we evaluated four time windows, as indicated in **C** and explained in Table S1. * indicates statistical significance, see Table S1. N = naïve, P = poorly trained, W = Well-trained.

**Figure S2.**
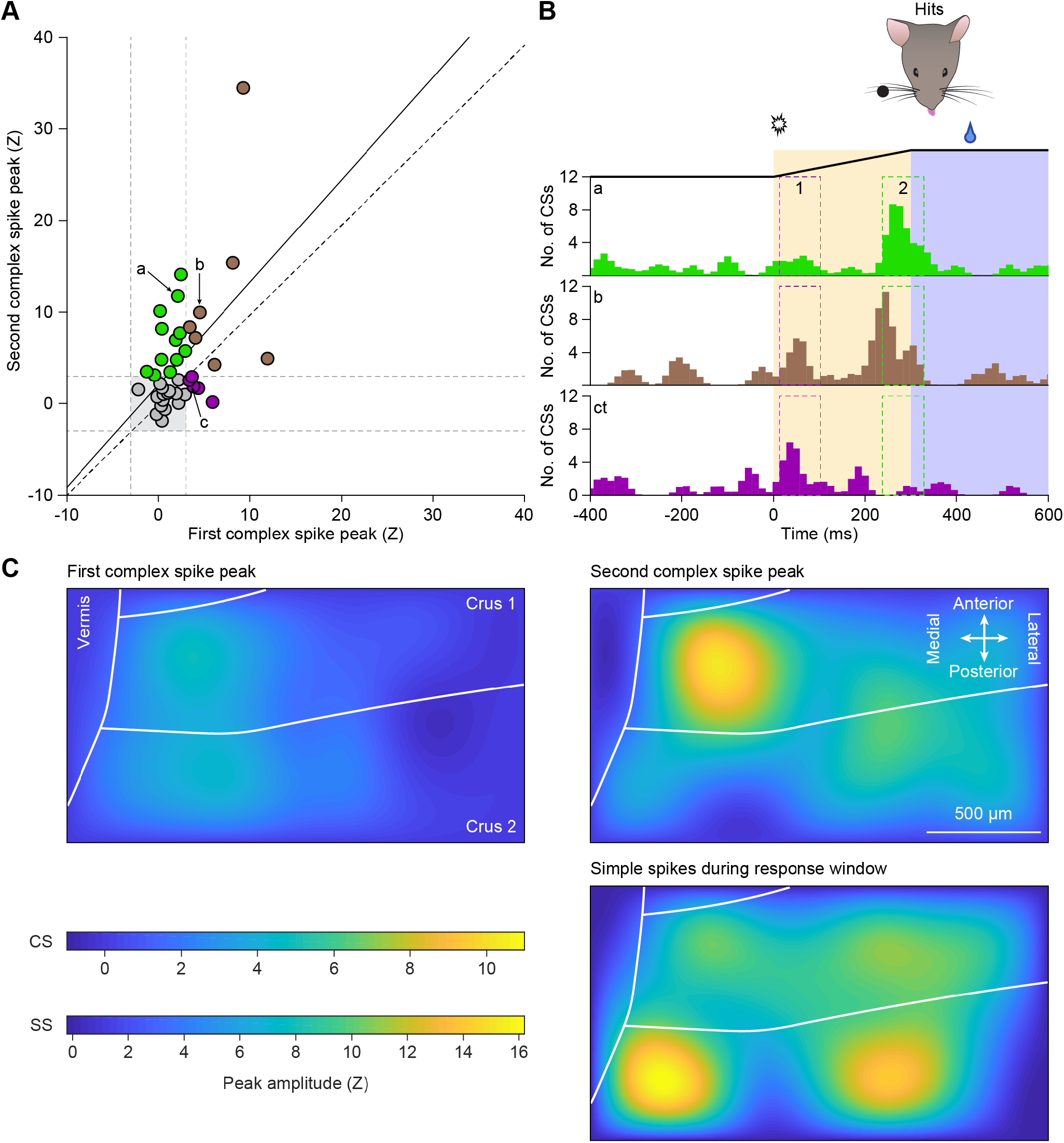
Spatiotemporal aspects of complex spikes and simple spikes, belonging to Fig. 3. **A**. At the level of individual Purkinje cells, the strength of the first (sound-evoked; 20-100 ms) and second (touch-induced; 240-320 ms) complex spike peak (see Fig. 3C) were weakly correlated. The “purple cells” preferentially fired during the first time window, the “green cells” during the second, and the “brown cells” during both. The solid line indicates the linear regression line (r = 0.35, *p* = 0.019, Spearman correlation test), the dotted line is at 45°, indicating equal strength of both peaks. **B**. Peri-stimulus time histograms of three example Purkinje cells. The numbers refer to their location in **A. C**. Relative strength of the first and second complex spike peak, respectively, as distributed over the area of crus 1 and crus 2, as well as that of the simple spike modulation during the response window.

**Figure S3.**
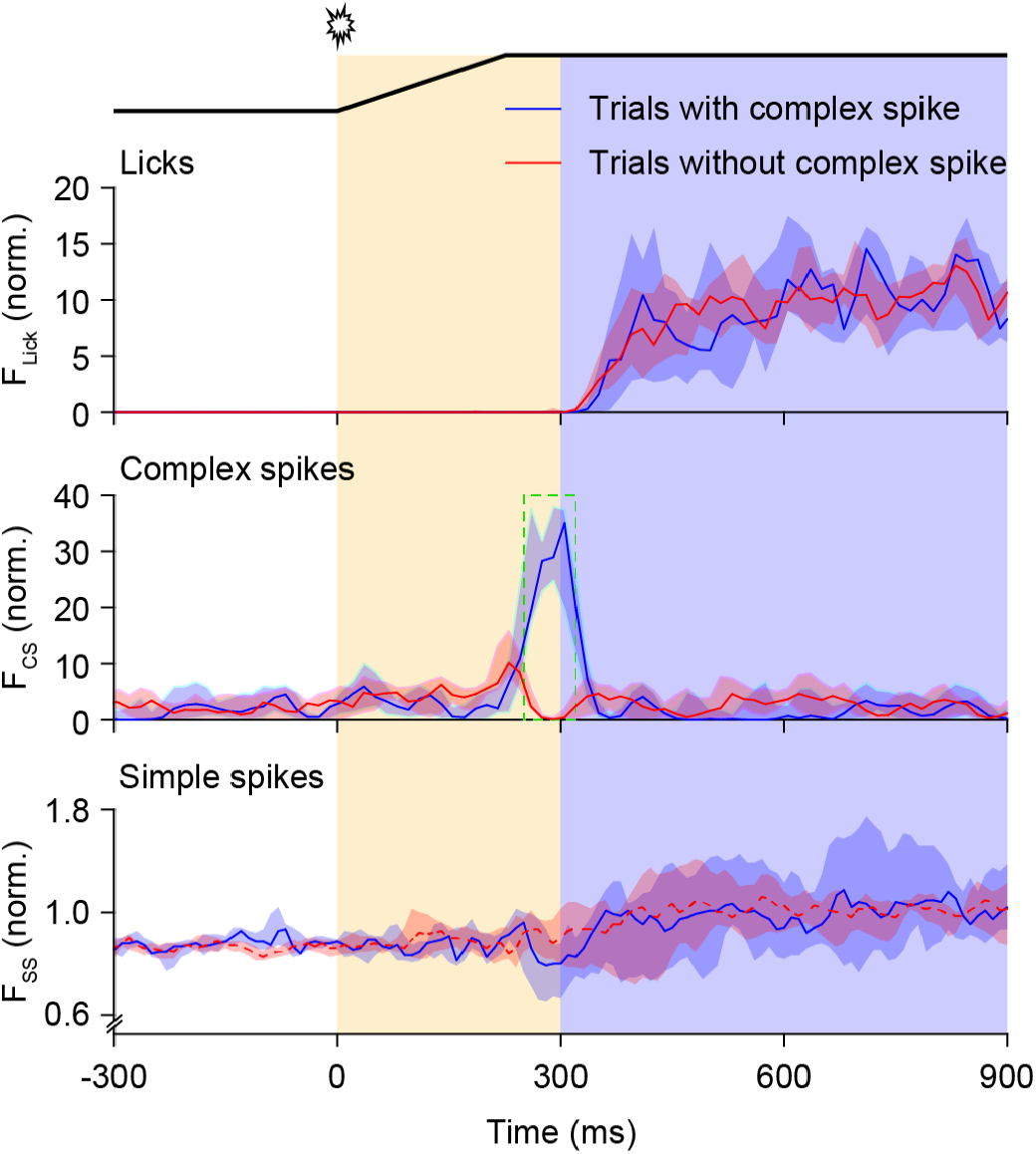
Complex spikes did not affect the licking behavior, belonging to Fig. 5. Comparing the trials during which a complex spike was fired during the second window of opportunity (240-320 ms after trial start; see Fig. 3C) with those trials that lacked a complex spike in that interval did not reveal any obvious difference in licking behavior.

**Figure S4.**
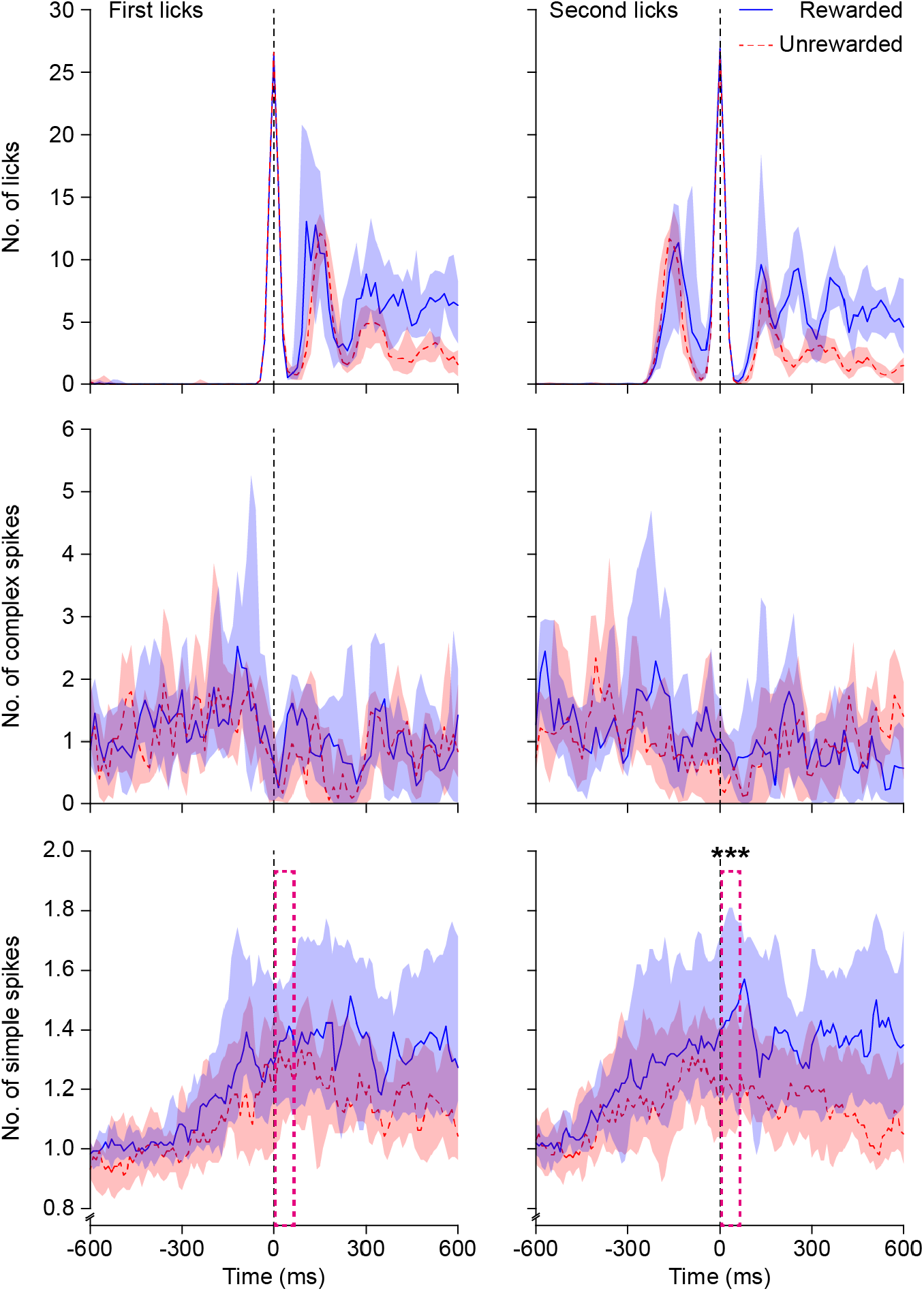
Effect of reward delivery on simple spike firing, belonging to Fig. 5. Peri-stimulus histograms of licks, complex spikes and simple spikes triggered on the first (left) or second (right) lick of bouts that were rewarded (blue) or unrewarded (red). During rewarded bouts, the first lick triggered a water reward. Rewarded lick bouts lasted longer than unrewarded ones. Note that the simple spikes after the second lick – thus at the moment that the mouse noticed that it got a reward or not – differed between rewarded and unrewarded licks (5-65 ms after detection of second lick: *p* = 0.007, W = 129, *n =* 19 Purkinje cells, Wilcoxon matched-pairs test).

**Figure S5.**
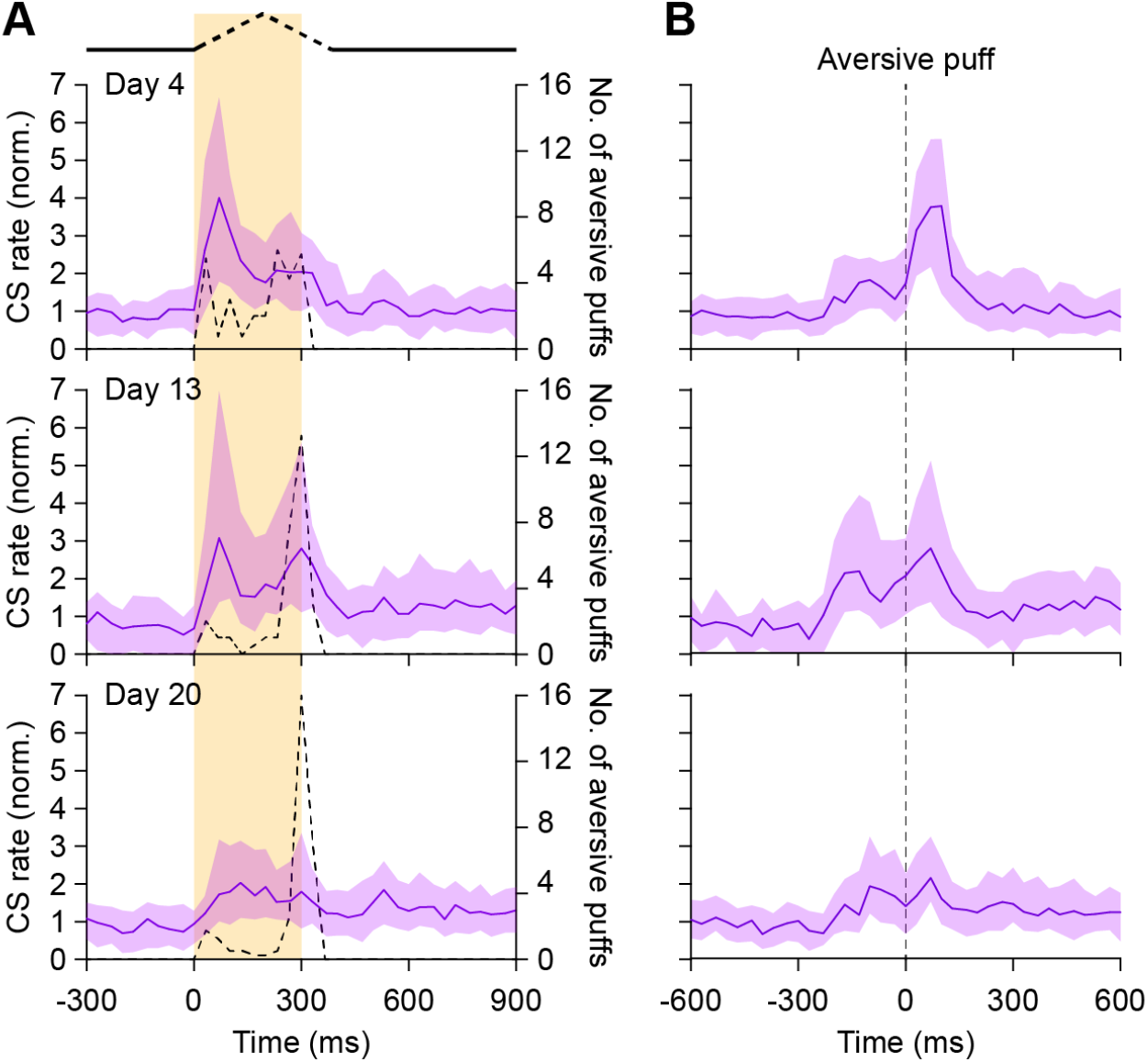
Impact of aversive puff diminishes with training, belonging to Fig. 6. **A**. When a mouse licked during the no-lick period of 300 ms following trial start, it received an aversive air puff to its nose, and the trial was aborted without the option to get a water reward. The dotted lines indicate the histogram of the occurrences of aversive puffs. Note that the aversive puffs were only applied during the no-lick period, but are indicated here with the same temporal resolution as the calcium imaging (30 Hz). Although early licking remained, the impact of the aversive puffs on complex spike firing (as measured with a miniscope, see Fig. 6) strongly diminished with time. **B**. This diminishing effect of the aversive puff on complex spike firing was further substantiated by triggering complex spike firing on the aversive puff. For both panels, only trials with aversive puffs were analyzed.

**Figure S6.**
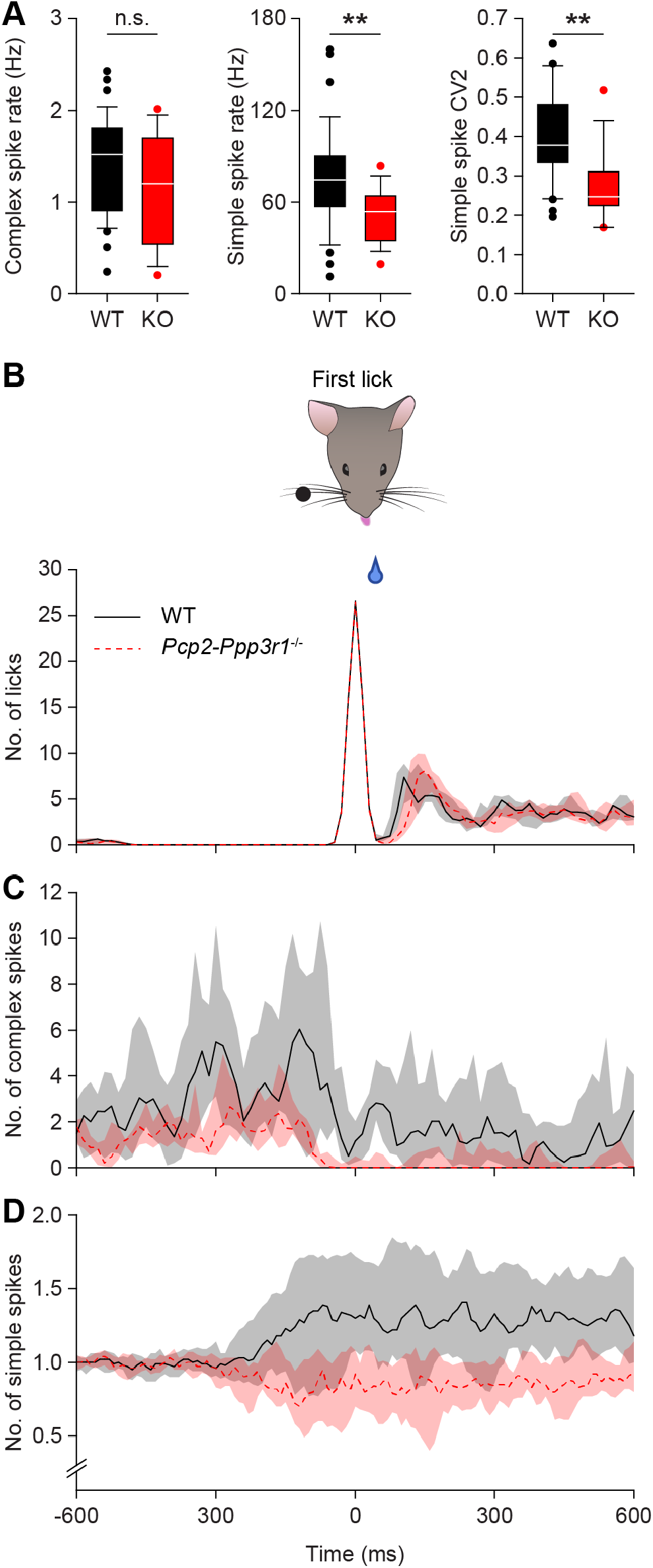
Purkinje cell responses during trials with licks in *Pcp2-Ppp3r1* KO mice, belonging to Fig. 7. **A**. Comparison of the average complex spike (left) and simple spike rate (middle) and simple spike CV2 in wild type and *Pcp2-Ppp3r1* KO mice. For statistics, see Table S2. Licks (**B**), complex spikes (**C**) and simple spikes (**D**) triggered on the first lick of bouts within the response window of hit trials. Note the decrease in complex spikes around licking start, as well as the suppressed simple spike firing during licking in *Pcp2-Ppp3r1* KO mice.

**Table S1.**
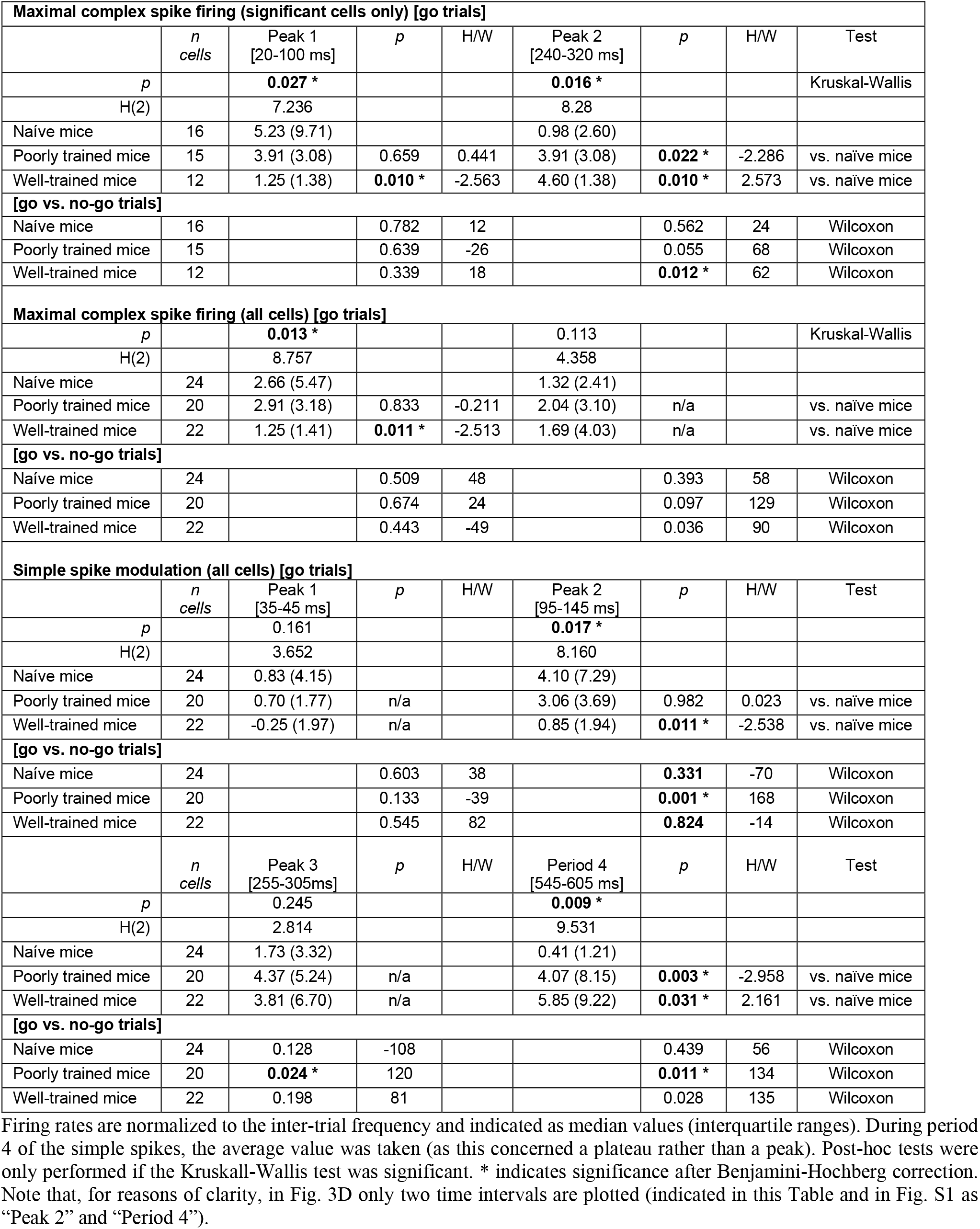
Statistical evaluation of complex spike and simple spike rates, belonging to Figs. 3 and S1.

| **Maximal complex spike firing (significant cells only) [go trials]** |  |  |  |  |  |  |  |  |
| --- | --- | --- | --- | --- | --- | --- | --- | --- |
|  | *n*<br>cells | Peak 1<br>[20-100 ms] | *p* | H/W | Peak 2<br>[240-320 ms] | *p* | H/W | Test |
| *p* |  | **0.027 \*** |  |  | **0.016 \*** |  |  | Kruskal-Wallis |
| H(2) |  | 7.236 |  |  | 8.28 |  |  |  |
| Naïve mice | 16 | 5.23 (9.71) |  |  | 0.98 (2.60) |  |  |  |
| Poorly trained mice | 15 | 3.91 (3.08) | 0.659 | 0.441 | 3.91 (3.08) | **0.022 \*** | -2.286 | vs. naïve mice |
| Well-trained mice | 12 | 1.25 (1.38) | **0.010 \*** | -2.563 | 4.60 (1.38) | **0.010 \*** | 2.573 | vs. naïve mice |
| **[go vs. no-go trials]** |  |  |  |  |  |  |  |  |
| Naïve mice | 16 |  | 0.782 | 12 |  | 0.562 | 24 | Wilcoxon |
| Poorly trained mice | 15 |  | 0.639 | -26 |  | 0.055 | 68 | Wilcoxon |
| Well-trained mice | 12 |  | 0.339 | 18 |  | **0.012 \*** | 62 | Wilcoxon |
| **Maximal complex spike firing (all cells) [go trials]** |  |  |  |  |  |  |  |  |
| *p* |  | **0.013 \*** |  |  | 0.113 |  |  | Kruskal-Wallis |
| H(2) |  | 8.757 |  |  | 4.358 |  |  |  |
| Naïve mice | 24 | 2.66 (5.47) |  |  | 1.32 (2.41) |  |  |  |
| Poorly trained mice | 20 | 2.91 (3.18) | 0.833 | -0.211 | 2.04 (3.10) | n/a |  | vs. naïve mice |
| Well-trained mice | 22 | 1.25 (1.41) | **0.011 \*** | -2.513 | 1.69 (4.03) | n/a |  | vs. naïve mice |
| **[go vs. no-go trials]** |  |  |  |  |  |  |  |  |
| Naïve mice | 24 |  | 0.509 | 48 |  | 0.393 | 58 | Wilcoxon |
| Poorly trained mice | 20 |  | 0.674 | 24 |  | 0.097 | 129 | Wilcoxon |
| Well-trained mice | 22 |  | 0.443 | -49 |  | 0.036 | 90 | Wilcoxon |
| **Simple spike modulation (all cells) [go trials]** |  |  |  |  |  |  |  |  |
|  | *n*<br>cells | Peak 1<br>[35-45 ms] | *p* | H/W | Peak 2<br>[95-145 ms] | *p* | H/W | Test |
| *p* |  | 0.161 |  |  | **0.017 \*** |  |  |  |
| H(2) |  | 3.652 |  |  | 8.160 |  |  |  |
| Naïve mice | 24 | 0.83 (4.15) |  |  | 4.10 (7.29) |  |  |  |
| Poorly trained mice | 20 | 0.70 (1.77) | n/a |  | 3.06 (3.69) | 0.982 | 0.023 | vs. naïve mice |
| Well-trained mice | 22 | -0.25 (1.97) | n/a |  | 0.85 (1.94) | **0.011 \*** | -2.538 | vs. naïve mice |
| **[go vs. no-go trials]** |  |  |  |  |  |  |  |  |
| Naïve mice | 24 |  | 0.603 | 38 |  | **0.331** | -70 | Wilcoxon |
| Poorly trained mice | 20 |  | 0.133 | -39 |  | **0.001 \*** | 168 | Wilcoxon |
| Well-trained mice | 22 |  | 0.545 | 82 |  | **0.824** | -14 | Wilcoxon |
|  | *n*<br>cells | Peak 3<br>[255-305ms] | *p* | H/W | Period 4<br>[545-605 ms] | *p* | H/W | Test |
| *p* |  | 0.245 |  |  | **0.009 \*** |  |  |  |
| H(2) |  | 2.814 |  |  | 9.531 |  |  |  |
| Naïve mice | 24 | 1.73 (3.32) |  |  | 0.41 (1.21) |  |  |  |
| Poorly trained mice | 20 | 4.37 (5.24) | n/a |  | 4.07 (8.15) | **0.003 \*** | -2.958 | vs. naïve mice |
| Well-trained mice | 22 | 3.81 (6.70) | n/a |  | 5.85 (9.22) | **0.031 \*** | 2.161 | vs. naïve mice |
| **[go vs. no-go trials]** |  |  |  |  |  |  |  |  |
| Naïve mice | 24 | 0.128 | -108 |  |  | 0.439 | 56 | Wilcoxon |
| Poorly trained mice | 20 | **0.024 \*** | 120 |  |  | **0.011 \*** | 134 | Wilcoxon |
| Well-trained mice | 22 | 0.198 | 81 |  |  | 0.028 | 135 | Wilcoxon |

**Table S2.** Spiking parameters of *Pcp2-Ppp3r1* KO mice, belonging to Fig. S6A.

|  | WT<br>(n = 37) | *Pcp2-Ppp3r1* KO<br>(n = 19) | *p* | U | Sign? | Test |
| --- | --- | --- | --- | --- | --- | --- |
| Complex spike rate | 1.5 (0.8) Hz | 1.2 (1.0) Hz | 0.081 | 250 | no | Mann-Whitney |
| Simple spike rate | 74.6 (30.37) Hz | 53.64 (26.3) Hz | **0.004** | 184 | Yes | Mann-Whitney |
| Simple spike CV2 | 0.38 ± 0.14 | 0.25 ± 0.08 | **<0.001** | 151 | Yes | Mann-Whitney |

Frequencies and CV2 are indicated as median values (interquartile ranges). Statistical significance (yes or no) is indicated after Bonferroni correction for multiple comparisons.

## References

Abdelgabar, A.R., Suttrup, J., Broersen, R., Bhandari, R., Picard, S., Keysers, C., De Zeeuw, C.I., and Gazzola, V. (2019). Action perception recruits the cerebellum and is impaired in patients with spinocerebellar ataxia. Brain 142, 3791–3805.

Albus, J.S. (1971). Theory of cerebellar function. Math Biosci 10, 25–61.

Allen, G., Buxton, R.B., Wong, E.C., and Courchesne, E. (1997). Attentional activation of the cerebellum independent of motor involvement. Science 275, 1940–1943.

Barski, J.J., Dethleffsen, K., and Meyer, M. (2000). Cre recombinase expression in cerebellar Purkinje cells. Genesis 28, 93–98.

Borke, R.C., Nau, M.E., and Ringler, R.L., Jr. (1983). Brain stem afferents of hypoglossal neurons in the rat. Brain Res 269, 47–55.

Bosman, L.W.J., Houweling, A.R., Owens, C.B., Tanke, N., Shevchouk, O.T., Rahmati, N., Teunissen, W.H.T., Ju, C., Gong, W., Koekkoek, S.K.E., and De Zeeuw, C.I. (2011). Anatomical pathways involved in generating and sensing rhythmic whisker movements. Front Integr Neurosci 5, 53.

Bosman, L.W.J., Koekkoek, S.K.E., Shapiro, J., Rijken, B.F.M., Zandstra, F., van der Ende, B., Owens, C.B., Potters, J.W., de Gruijl, J.R., Ruigrok, T.J.H., and De Zeeuw, C.I. (2010). Encoding of whisker input by cerebellar Purkinje cells. J Physiol 588, 3757–3783.

Boyd, C.A.R. (2010). Cerebellar agenesis revisited. Brain 133, 941–944.

Brissenden, J.A., Tobyne, S.M., Osher, D.E., Levin, E.J., Halko, M.A., and Somers, D.C. (2018). Topographic cortico-cerebellar networks revealed by visual attention and working memory. Curr Biol 28, 3364–3372 e3365.

Brooks, J.X., Carriot, J., and Cullen, K.E. (2015). Learning to expect the unexpected: rapid updating in primate cerebellum during voluntary self-motion. Nat Neurosci 18, 1310–1317.

Bryant, J.L., Boughter, J.D., Gong, S., Ledoux, M.S., and Heck, D.H. (2010). Cerebellar cortical output encodes temporal aspects of rhythmic licking movements and is necessary for normal licking frequency. Eur J Neurosci 32, 41–52.

Buschman, T.J., and Kastner, S. (2015). From behavior to neural dynamics: An integrated theory of attention. Neuron 88, 127–144.

Cant, N.B., and Oliver, D.L. (2018). Overview of auditory projection pathways and intrinsic microcircuits. In The mammalian auditory pathways, D. Oliver, N. Cant, R. Fay, and A. Popper, eds. (Cham: Springer), pp. 7–39.

Castellucci, V., Pinsker, H., Kupfermann, I., and Kandel, E.R. (1970). Neuronal mechanisms of habituation and dishabituation of the gill-withdrawal reflex in Aplysia. Science 167, 1745–1748.

Cayco-Gajic, N.A., and Silver, R.A. (2019). Re-evaluating circuit mechanisms underlying pattern separation. Neuron 101, 584–602.

Chabrol, F.P., Blot, A., and Mrsic-Flogel, T.D. (2019). Cerebellar contribution to preparatory activity in motor neocortex. Neuron 103, 506–519 e504.

Clutton-Brock, T.H., O’Riain, M.J., Brotherton, P.N.M., Gaynor, D., Kansky, R., Griffin, A.S., and Manser, M. (1999). Selfish sentinels in cooperative mammals. Science 284, 1640–1644.

Coesmans, M., Weber, J.T., De Zeeuw, C.I., and Hansel, C. (2004). Bidirectional parallel fiber plasticity in the cerebellum under climbing fiber control. Neuron 44, 691–700.

de Groot, A., van den Boom, B.J.G., van Genderen, R.M., Coppens, J., van Veldhuijzen, J., Bos, J., Hoedemaker, H., Negrello, M., Willuhn, I., De Zeeuw, C.I., and Hoogland, T.M. (2020). NINscope, a versatile miniscope for multi-region circuit investigations. Elife 9.

De Zeeuw, C.I. (2020). Bidirectional learning in upbound and downbound microzones of the cerebellum. Nat Rev Neurosci in press.

De Zeeuw, C.I., Hoebeek, F.E., Bosman, L.W.J., Schonewille, M., Witter, L., and Koekkoek, S.K. (2011). Spatiotemporal firing patterns in the cerebellum. Nat Rev Neurosci 12, 327–344.

De Zeeuw, C.I., and Ten Brinke, M.M. (2015). Motor learning and the cerebellum. Cold Spring Harb Perspect Biol 7, a021683.

Deverett, B., Koay, S.A., Oostland, M., and Wang, S.S. (2018). Cerebellar involvement in an evidence-accumulation decision-making task. Elife 7.

Frost, W.N., Castellucci, V.F., Hawkins, R.D., and Kandel, E.R. (1985). Monosynaptic connections made by the sensory neurons of the gill- and siphon-withdrawal reflex in *Aplysia* participate in the storage of long-term memory for sensitization. Proc Natl Acad Sci U S A 82, 8266–8269.

Gao, Z., Davis, C., Thomas, A.M., Economo, M.N., Abrego, A.M., Svoboda, K., De Zeeuw, C.I., and Li, N. (2018). A cortico-cerebellar loop for motor planning. Nature 563, 113–116.

Gao, Z., Van Beugen, B.J., and De Zeeuw, C.I. (2012). Distributed synergistic plasticity and cerebellar learning. Nat Rev Neurosci 13, 619–635.

Heffley, W., and Hull, C. (2019). Classical conditioning drives learned reward prediction signals in climbing fibers across the lateral cerebellum. Elife 8.

Heffley, W., Song, E.Y., Xu, Z., Taylor, B.N., Hughes, M.A., McKinney, A., Joshua, M., and Hull, C. (2018). Coordinated cerebellar climbing fiber activity signals learned sensorimotor predictions. Nat Neurosci 21, 1431–1441.

Helmuth, L.L., Ivry, R.B., and Shimizu, N. (1997). Preserved performance by cerebellar patients on tests of word generation, discrimination learning, and attention. Learn Mem 3, 456–474.

Herzfeld, D.J., Kojima, Y., Soetedjo, R., and Shadmehr, R. (2018). Encoding of error and learning to correct that error by the Purkinje cells of the cerebellum. Nat Neurosci 21, 736–743.

Hull, C. (2020). Prediction signals in the cerebellum: beyond supervised motor learning. Elife 9.

Ito, M. (2000). Mechanisms of motor learning in the cerebellum. Brain Res 886, 237–245.

Ito, M. (2002). The molecular organization of cerebellar long-term depression. Nat Rev Neurosci 3, 896–902.

Ito, M., Yoshida, M., and Obata, K. (1964). Monosynaptic inhibition of the intracerebellar nuclei induced rom the cerebellar cortex. Experientia 20, 575–576.

Ivry, R.B., and Keele, S.W. (1989). Timing functions of the cerebellum. Journal of cognitive neuroscience 1, 136–152.

Jirenhed, D.A., Bengtsson, F., and Hesslow, G. (2007). Acquisition, extinction, and reacquisition of a cerebellar cortical memory trace. J Neurosci 27, 2493–2502.

Ju, C., Bosman, L.W.J., Hoogland, T.M., Velauthapillai, A., Murugesan, P., Warnaar, P., Negrello, M., and De Zeeuw, C.I. (2019). Neurons of the inferior olive respond to broad classes of sensory input while subject to homeostatic control. J Physiol 597, 2483–2514.

Knudsen, E.I. (2018). Neural circuits that mediate selective attention: A comparative perspective. Trends Neurosci 41, 789–805.

Koekkoek, S.K.E., Hulscher, H.C., Dortland, B.R., Hensbroek, R.A., Elgersma, Y., Ruigrok, T.J.H., and De Zeeuw, C.I. (2003). Cerebellar LTD and learning-dependent timing of conditioned eyelid responses. Science 301, 1736–1739.

Kostadinov, D., Beau, M., Pozo, M.B., and Häusser, M. (2019). Predictive and reactive reward signals conveyed by climbing fiber inputs to cerebellar Purkinje cells. Nat Neurosci 22, 950–962.

Kuhn, B., Ozden, I., Lampi, Y., Hasan, M.T., and Wang, S.S.H. (2012). An amplified promoter system for targeted expression of calcium indicator proteins in the cerebellar cortex. Frontiers in neural circuits 6, 49.

Larry, N., Yarkoni, M., Lixenberg, A., and Joshua, M. (2019). Cerebellar climbing fibers encode expected reward size. Elife 8.

Le, T.H., Pardo, J.V., and Hu, X. (1998). 4 T-fMRI study of nonspatial shifting of selective attention: cerebellar and parietal contributions. J Neurophysiol 79, 1535–1548.

Lowe, A.A. (1980). The neural regulation of tongue movements. Prog Neurobiol 15, 295–344.

Marr, D. (1969). A theory of cerebellar cortex. J Physiol 202, 437–470.

McCormick, D.A., and Thompson, R.F. (1984). Cerebellum: essential involvement in the classically conditioned eyelid response. Science 223, 296–299.

McElvain, L.E., Friedman, B., Karten, H.J., Svoboda, K., Wang, F., Deschênes, M., and Kleinfeld, D. (2018). Circuits in the rodent brainstem that control whisking in concert with other orofacial motor actions. Neuroscience 368, 152–170.

Moberget, T., and Ivry, R.B. (2019). Prediction, psychosis, and the cerebellum. Biol Psychiatry Cogn Neurosci Neuroimaging 4, 820–831.

Moore, D.R. (1991). Anatomy and physiology of binaural hearing. Audiology 30, 125–134.

Negrello, M., Warnaar, P., Romano, V., Owens, C.B., Lindeman, S., Iavarona, E., Spanke, J.K., Bosman, L.W.J., and De Zeeuw, C.I. (2019). Quasiperiodic rhythms of the inferior olive. PLoS Comput Biol 15, e1006475.

Ohmae, S., and Medina, J.F. (2015). Climbing fibers encode a temporal-difference prediction error during cerebellar learning in mice. Nat Neurosci 18, 1798–1803.

Ohtsuki, G., Piochon, C., and Hansel, C. (2009). Climbing fiber signaling and cerebellar gain control. Front Cell Neurosci 3, 4.

Pnevmatikakis, E.A., and Giovannucci, A. (2017). NoRMCorre: An online algorithm for piecewise rigid motion correction of calcium imaging data. J Neurosci Methods 291, 83–94.

Rahmati, N., Owens, C.B., Bosman, L.W.J., Spanke, J.K., Lindeman, S., Gong, W., Potters, J.W., Romano, V., Voges, K., Moscato, L., et al. (2014). Cerebellar potentiation and learning a whisker-based object localization task with a time response window. J Neurosci 34, 1949–1962.

Romano, V., De Propris, L., Bosman, L.W.J., Warnaar, P., ten Brinke, M.M., Lindeman, S., Ju, C., Velauthapillai, A., Spanke, J.K., Middendorp Guerra, E., et al. (2018). Potentiation of cerebellar Purkinje cells facilitates whisker reflex adaptation through increased simple spike activity. eLife 7, e38852.

Romano, V., Reddington, A.L., Cazzanelli, S., Mazza, R., Ma, Y., Strydis, C., Negrello, M., Bosman, L.W.J., and De Zeeuw, C.I. (2020). Functional convergence of autonomic and sensorimotor processing in the lateral cerebellum. Cell Rep 32, 107867.

Santema, P., and Clutton-Brock, T. (2013). Meerkat helpers increase sentinel behaviour and bipedal vigilance in the presence of pups. Animal Behaviour 85, 655–661.

Sauerbrei, B.A., Lubenov, E.V., and Siapas, A.G. (2015). Structured variability in Purkinje cell activity during locomotion. Neuron 87, 840–852.

Schonewille, M., Belmeguenai, A., Koekkoek, S.K., Houtman, S.H., Boele, H.J., van Beugen, B.J., Gao, Z., Badura, A., Ohtsuki, G., Amerika, W.E., et al. (2010). Purkinje cell-specific knockout of the protein phosphatase PP2B impairs potentiation and cerebellar motor learning. Neuron 67, 618–628.

Sendhilnathan, N., Ipata, A.E., and Goldberg, M.E. (2020). Neural correlates of reinforcement learning in mid-lateral cerebellum. Neuron 106, 188–198 e185.

Smith, E.E., and Jonides, J. (1999). Storage and executive processes in the frontal lobes. Science 283, 1657–1661.

Steinmetz, J.E., Logan, C.G., Rosen, D.J., Thompson, J.K., Lavond, D.G., and Thompson, R.F. (1987). Initial localization of the acoustic conditioned stimulus projection system to the cerebellum essential for classical eyelid conditioning. Proc Natl Acad Sci U S A 84, 3531–3535.

Suvrathan, A., Payne, H.L., and Raymond, J.L. (2016). Timing rules for synaptic plasticity matched to behavioral function. Neuron 92, 959–967.

Ten Brinke, M.M., Boele, H.J., and De Zeeuw, C.I. (2019). Conditioned climbing fiber responses in cerebellar cortex and nuclei. Neurosci Lett 688, 26–36.

Ten Brinke, M.M., Boele, H.J., Spanke, J.K., Potters, J.W., Kornysheva, K., Wulff, P., IJpelaar, A.C.H.G., Koekkoek, S.K.E., and De Zeeuw, C.I. (2015). Evolving models of Pavlovian conditioning: cerebellar cortical dynamics in awake behaving mice. Cell Rep 13, 1977–1988.

Teune, T.M., van der Burg, J., van der Moer, J., Voogd, J., and Ruigrok, T.J. (2000). Topography of cerebellar nuclear projections to the brain stem in the rat. Progress in brain research 124, 141–172.

Thach, W.T., Jr. (1967). Somatosensory receptive fields of single units in cat cerebellar cortex. J Neurophysiol 30, 675–696.

Tsutsumi, S., Hidaka, N., Isomura, Y., Matsuzaki, M., Sakimura, K., Kano, M., and Kitamura, K. (2019). Modular organization of cerebellar climbing fiber inputs during goal-directed behavior. Elife 8.

Tzvi, E., Koeth, F., Karabanov, A.N., Siebner, H.R., and Krämer, U.M. (2020). Cerebellar – Premotor cortex interactions underlying visuomotor adaptation. NeuroImage 220, 117142.

Vinueza Veloz, M.F., Zhou, K., Bosman, L.W.J., Potters, J.W., Negrello, M., Seepers, R.M., Strydis, C., Koekkoek, S.K.E., and De Zeeuw, C.I. (2015). Cerebellar control of gait and interlimb coordination. Brain structure & function 220, 3513–3536.

Voges, K., Wu, B., Post, L., Schonewille, M., and De Zeeuw, C.I. (2017). Mechanisms underlying vestibulo-cerebellar motor learning in mice depend on movement direction. J Physiol 595, 5301–5326.

Wagner, M.J., Kim, T.H., Savall, J., Schnitzer, M.J., and Luo, L.Q. (2017). Cerebellar granule cells encode the expectation of reward. Nature 544, 96–100.

Witter, L., Canto, C.B., Hoogland, T.M., de Gruijl, J.R., and De Zeeuw, C.I. (2013). Strength and timing of motor responses mediated by rebound firing in the cerebellar nuclei after Purkinje cell activation. Frontiers in neural circuits 7, 133.

Wolpert, D.M., Miall, R.C., and Kawato, M. (1998). Internal models in the cerebellum. Trends Cogn Sci 2, 338–347.

Yang, Y., and Lisberger, S.G. (2014). Purkinje-cell plasticity and cerebellar motor learning are graded by complex-spike duration. Nature 510, 529–532.

Yu, C., Derdikman, D., Haidarliu, S., and Ahissar, E. (2006). Parallel thalamic pathways for whisking and touch signals in the rat. PLoS Biol 4, e124.

Zeng, H., Chattarji, S., Barbarosie, M., Rondi-Reig, L., Philpot, B.D., Miyakawa, T., Bear, M.F., and Tonegawa, S. (2001). Forebrain-specific calcineurin knockout selectively impairs bidirectional synaptic plasticity and working/episodic-like memory. Cell 107, 617–629.

Zhou, H., Lin, Z., Voges, K., Ju, C., Gao, Z., Bosman, L.W.J., Ruigrok, T.J.H., Hoebeek, F.E., De Zeeuw, C.I., and Schonewille, M. (2014). Cerebellar modules operate at different frequencies. eLife 3, e02536.

Zhou, P., Resendez, S.L., Rodriguez-Romaguera, J., Jimenez, J.C., Neufeld, S.Q., Giovannucci, A., Friedrich, J., Pnevmatikakis, E.A., Stuber, G.D., Hen, R., et al. (2018). Efficient and accurate extraction of in vivo calcium signals from microendoscopic video data. Elife 7.

